# Local translation of yeast *ERG4* mRNA at the endoplasmic reticulum requires the brefeldin A resistance protein Bfr1

**DOI:** 10.1101/656405

**Authors:** Srinivas Manchalu, Nitish Mittal, Anne Spang, Ralf-Peter Jansen

**Affiliations:** Interfaculty Institute of Biochemistry, University of Tübingen, Tübingen, Germany; Biozentrum, University of Basel, Basel, Switzerland

**Keywords:** RNA-binding protein, yeast, endoplasmic reticulum, Brefeldin A resistance, ribosomal occupancy, RNA-protein interaction

## Abstract

Brefeldin A resistance factor 1 (Bfr1p) is a non-essential RNA-binding protein and multi-copy suppressor of brefeldin A sensitivity in *Saccharomyces cerevisiae*. Deletion of *BFR1* leads to multiple defects, including altered cell shape and size, change in ploidy, induction of P-bodies and chromosomal mis-segregation. Bfr1p has been shown to associate with polysomes, binds to several hundred mRNAs, and can target some of them to P-bodies. Although this implies a role of Bfr1p in translational control of mRNAs, its molecular function remains elusive. In the present study, we show that mutations in RNA-binding residues of Bfr1p impede its RNA-dependent co-localization with ER, yet do not mimic the known cellular defects seen upon *BFR1* deletion. However, a Bfr1 RNA-binding mutant is impaired in binding to *ERG4* mRNA which encodes an enzyme required for the final step of ergosterol biosynthesis. Consistently, *bfr1*Δ strains show a strong reduction in Erg4p protein levels. Polysome profiling of *bfr1*Δ or *bfr1* mutant strains reveals a strong shift of *ERG4* mRNA to polysomes, consistent with a function of Bfr1p in elongation. Collectively, our data reveal that Bfr1 has at least two separable functions: one in RNA-binding and elongation during translation, in particular at the ER membrane, and one in ploidy control or chromosome segregation.

## INTRODUCTION

Function and lifetime of mRNAs are largely controlled by RNA-binding proteins (RBPs) that can regulate RNA folding, stability, translation, localization or access by other proteins (Neriec and Percipalle 2018; Björk and Wieslander 2017; Lunde et al. 2007; Dreyfuss et al. 2002). These proteins can bind RNA co-transcriptionally at birth and dynamically associate/dissociate throughout the lifetime of mRNAs. RNA recognition and binding generally occurs via RNA-binding domains (RBDs). Common RBDs include the RNA recognition motif (RRM; Oubridge et al. 1994), the heterologous nuclear RNA-binding protein K homology (KH) domain (Lewis et al. 2000), the double-stranded RNA binding domain (dsRBD; Ryter and Schultz 1998), or zinc-finger domains (Lu et al. 2003); for a detailed and more complete list, see (Auweter et al. 2006; Lunde et al. 2007). However, high-throughput RNA-binding studies have revealed a large number of RBPs lacking known RBDs, implicating the presence of even more RBPs than previously anticipated (Baltz et al. 2012; Castello et al. 2012; Kwon et al. 2013; Beckmann et al. 2015; Castello et al. 2015; Hentze et al. 2018). RNA binding of these proteins frequently occurs via intrinsically disordered regions (IDRs; (Hentze et al. 2018)) or nucleotide-binding sites (Castello et al. 2016; 2015). Many of these proteins like some enzymes have functions additional to RNA binding. Furthermore, in many cases we do not know the biological function of RNA binding or even their RNA targets.

The 55 kDa budding yeast Bfr1 protein is a non-essential RBP lacking common RBDs. The gene had been originally identified in a screen for genes relieving the growth defect seen in yeast cells carrying a mutation in *SEC21* if exposed to the drug brefeldin A (Jackson and Képès 1994) and hence named brefeldin A resistance factor 1 (*BFR1*). Since brefeldin A inhibits transport out of the Golgi apparatus, this early work suggested that Bfr1p is involved in membrane trafficking (Jackson and Képès 1994). However, later studies revealed a role for Bfr1p in mRNA metabolism (Lang et al. 2001). Bfr1p co-fractionates with polysomes, implying a function in translation. In addition, it associates with RNA-protein particles (RNPs) that contain the mRNA-binding protein Scp160p (Lang and Fridovich-Keil 2000) and the ribosome-associated protein Asc1p (Sezen et al. 2009). Co-purification of Bfr1p with mRNPs is independent of Scp160p but Bfr1p is required for Scp160p’s association with polysomes (Lang et al. 2001). Consistent with its putative role in translation, Bfr1p is present in the cytoplasm but also localizes to the endoplasmic reticulum (ER), possibly due to its interaction with polyribosomes translating membrane or secreted proteins (Lang et al. 2001).

Potential RNA targets of Bfr1p have been revealed in several studies using RBP-immunoprecipitation coupled with microarray analysis (RIP-Chip; (Hogan et al. 2008)), crosslinking and immunoprecipitation (CLIP; (Mitchell et al. 2013)), or RNA-tagging (Lapointe et al. 2015). All studies concurrently reported that Bfr1p binds to several hundred different mRNAs. The set of Bfr1p-associated mRNAs largely overlaps with those that encode proteins translated at the ER (Lapointe et al. 2015; Jan et al. 2014). A similar observation had been made for Scp160p (Hogan et al. 2008; Lapointe et al. 2015), supporting the idea that the protein is implicated in ER-based translation.

In addition, Bfr1p functions in stress-related mRNA decay. During glucose depletion, Bfr1p relocates to P-bodies at a late stage of P-body formation (Simpson et al. 2014). It is also required for targeting of mRNAs to P-bodies at a late phase of stress. In unstressed cells, Bfr1p as well as Scp160 act as negative regulators of P-body formation, probably by protecting mRNA at polysomes (Weidner et al. 2014). Lack of any of these proteins results in P-body like structures even under optimal growth conditions (Weidner et al. 2014). Loss of Bfr1p additionally leads to changes in the cell ploidy (Jackson and Képès 1994). Such ploidy changes could be the result of chromosome missegregation due to a failure to build a functional mitotic bipolar spindle. Monopolar spindles can arise from defects in spindle pole body (SPB) duplication and the SPB component Bbp1p has been reported to bind to Bfr1p (Xue et al. 1996). In addition, Bfr1p interacts with the SESA (Smy2-Eap1-Scp160-Asc1) network that controls spindle pole duplication (Sezen et al. 2009).

All previous studies that reported on phenotypes associated with loss of Bfr1p were performed with *BFR1* deletion cells. Since loss of Bfr1p results in alterations of the genome (Jackson and Képès 1994), it remained unclear, if the observed effects are a direct consequence of Bfr1p loss and to what extent loss of RNA binding or mis-regulation of Bfr1p targets contribute to the phenotypes. Therefore, we generated Bfr1p mutants with point mutations in known RNA-binding sites in order to establish a link between mRNA binding, brefeldin A resistance, ploidy control, and translation. Here, we demonstrate that RNA-binding is required for Bfr1p function in translational regulation of mRNAs including *ERG4* that encodes an ergosterol synthesizing enzyme, and that this function is independent of Bfr1’s role in ploidy maintenance.

## RESULTS

### RNA-binding of Bfr1p is required for its localization to the ER

The non-canonical RBP Bfr1p is involved in several cellular processes including transfer and retention of mRNAs to P-bodies (Simpson et al. 2014; Weidner et al. 2014), translation (Lang et al. 2001), and response to functional failure of the yeast spindle pole body or kinetochore (Sezen et al. 2009; Low et al. 2014). We generated *bfr1* mutants in order to address the question of whether RNA-binding is required for all processes. Since Bfr1p lacks canonical RBDs, we made alanine substitutions at amino acid positions that had previously been identified to cross-link with RNA *in-vivo* (Kramer et al. 2014; Lapointe et al. 2015) (Figure S1). Out of the six RNA contact sites have been described, we initially focused on the two amino acids (K138, F239) that are most highly conserved in Bfr1p among different yeast species (Figures 1A and S1). In addition, we generated a shorter Bfr1p variant that lacks the third coiled coil domain, which is devoid of any identified RNA contact sites (Kramer et al. 2014). When expressed as carboxyterminal GFP fusion proteins in a *bfr1*Δ yeast strain, Bfr1p mutant proteins with point mutations in two cross-link sites (K139 or F293) or in all six sites are expressed similarly (Figure S2A and S2B) as endogenous full-length Bfr1p (Bfr1p WT FL). Similar results were obtained for the deletion mutant Bfr1p(1-397). In exponentially growing yeast cells, Bfr1p is located in the cytoplasm and at the ER (Lang et al. 2001), which is recapitulated by our Bfr1-GFP fusion protein (Figure 1B and 1C). Whereas deletion of the third coiled coil domain does not affect this distribution, mutation of all six conserved RNA cross-link sites as well as also the single exchange of F239A results in a partial loss of Bfr1p from the ER as determined by live cell imaging and subcellular fractionation (Figure 1B and C). The majority of wild-type Bfr1p co-sediments with a membrane fraction containing ER as judged by the distribution of the ER marker Sec61p. Similarly, the signal of Bfr1p(1-397) in the membrane fraction is also more prominent than in the cytoplasmic fraction. In contrast, the ratio of Bfr1p between membrane and cytoplasmic fraction is inversed for the *bfr1mutF239A* mutant. Out of all the single point mutations, F239 is the only cross-link site for which a mutation affects ER localization. Single mutations in each of the other five sites result in no change in Bfr1p distribution (Figure S2C). In order to investigate if the loss of ER localization of *bfr1mut6* and *bfr1mutF293A* proteins is due to loss of RNA binding, we followed the distribution of wild-type Bfr1p in subcellular distribution in lysates treated with RNase A (Figure 1D). RNA degradation results in a similar redistribution of Bfr1p from the membrane fractions to a cytosolic fraction, same as that observed for the mutant proteins. This suggest that RNA-binding ability of Bfr1p is required for its localization to the ER in yeast.

**Figure 1.**
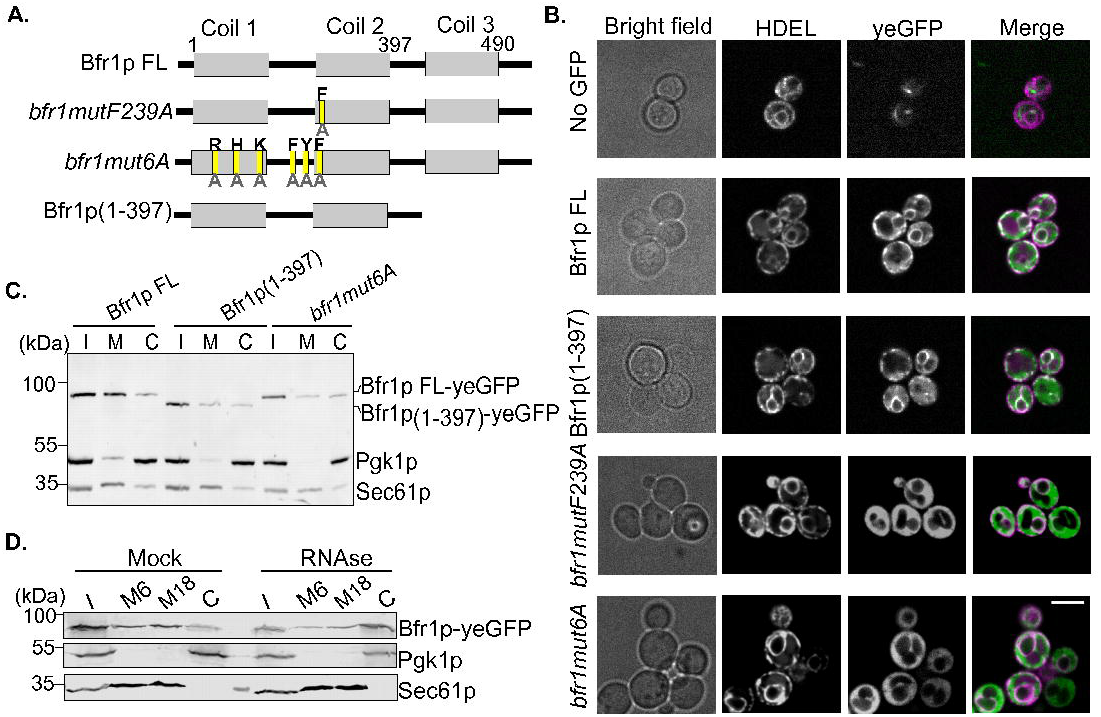
Mutations in RNA-binding residues of Bfr1p affects the Bfr1p-ER localization. A. Schematic view of Bfr1p constructs used in this study. All four constructs were fused to a C-terminal yeGFP. RNA binding residues R38, H79, K138, F211, Y225 and F239 were replaced with alanine and mutant *bfr1*mut*6A* contains all six mutations. Bfr1p (1-397) is a truncation lacking the third coiled coil. B. Intracellular distribution of yeGFP-tagged Bfr1p constructs. HDEL-DsRed serves as ER marker. Bfr1p variants of mutants *bfr1*mut*F239A* and *bfr1mut6A* show loss of co-localization between Bfr1p and the ER. Scale bar, 8 µm. C. Distribution of Bfr1p constructs in subcellular fractionation. I: input; M: membrane fraction (pellet from 18000 x*g*); C: cytoplasmic fraction (supernatant from 18000 x*g*). Pgk1p serves as cytosolic and Sec61p as ER markers. D. Redistribution of Bfr1p between membrane and cytosolic fractions upon RNAse treatment. M6: membrane fraction (pellet from 6000 x*g*), M18: membrane fraction (pellet from 18000 x*g*) and C: cytoplasmic fraction (supernatant from 18000 x*g*).

Since deletion of *BFR1* is associated with various phenotypes, we also tested if *bfr1mut6* or *bfr1mutF293A* affect ploidy maintenance (Jackson and Képès 1994; Xue et al. 1996), resistance to brefeldin A (Jackson and Képès 1994), or premature formation of P-bodies (Weidner et al. 2014). As described before, complete loss of *BFR1* results in a ploidy shift from 1N to 2N as exemplified by the shift of DNA peaks from 1C/2C (haploid) to 2C/4C (diploid) in flow cytometry analysis (Figure 2A). However, a similar shift is not seen in the tested mutants (Figure 2A). Overexpression of *BFR1* has been demonstrated to rescue the brefeldin A sensitivity of a yeast mutant lacking the *ERG6* gene (Graham et al. 1993; Shah and Klausner 1993). Since all tested *bfr1* point mutants including mutant *bfr1mut6A* are still able to rescue *erg6*Δ to a similar level as the wild type (Figure 2B), we conclude that none of these sites are important for suppression of the brefeldin A sensitivity in *erg6*Δ. Similarly, none of the RNA-binding mutants lead to occurrence of premature P-bodies is observed in *bfr1*Δ cells. Formation of P-bodies in wild type cells or *bfr1*Δ cells expressing various Bfr1p-GFP variants was assessed by co-expression of Edc3p-mCherry, a bona-fide P body marker (Weidner et al. 2014). Formation of Edc3p-positive cytoplasmic foci in wild type cells is induced by transferring cells into medium lacking glucose (Figure 2C, right panels), a potent inducer of P-bodies. Whereas *bfr1*Δ cells form Edc3p foci also in the presence of glucose, expression of full-length Bfr1p or Bfr1p RNA-binding mutants rescues this phenotype (Figure 2C, left panels).

**Figure 2.**
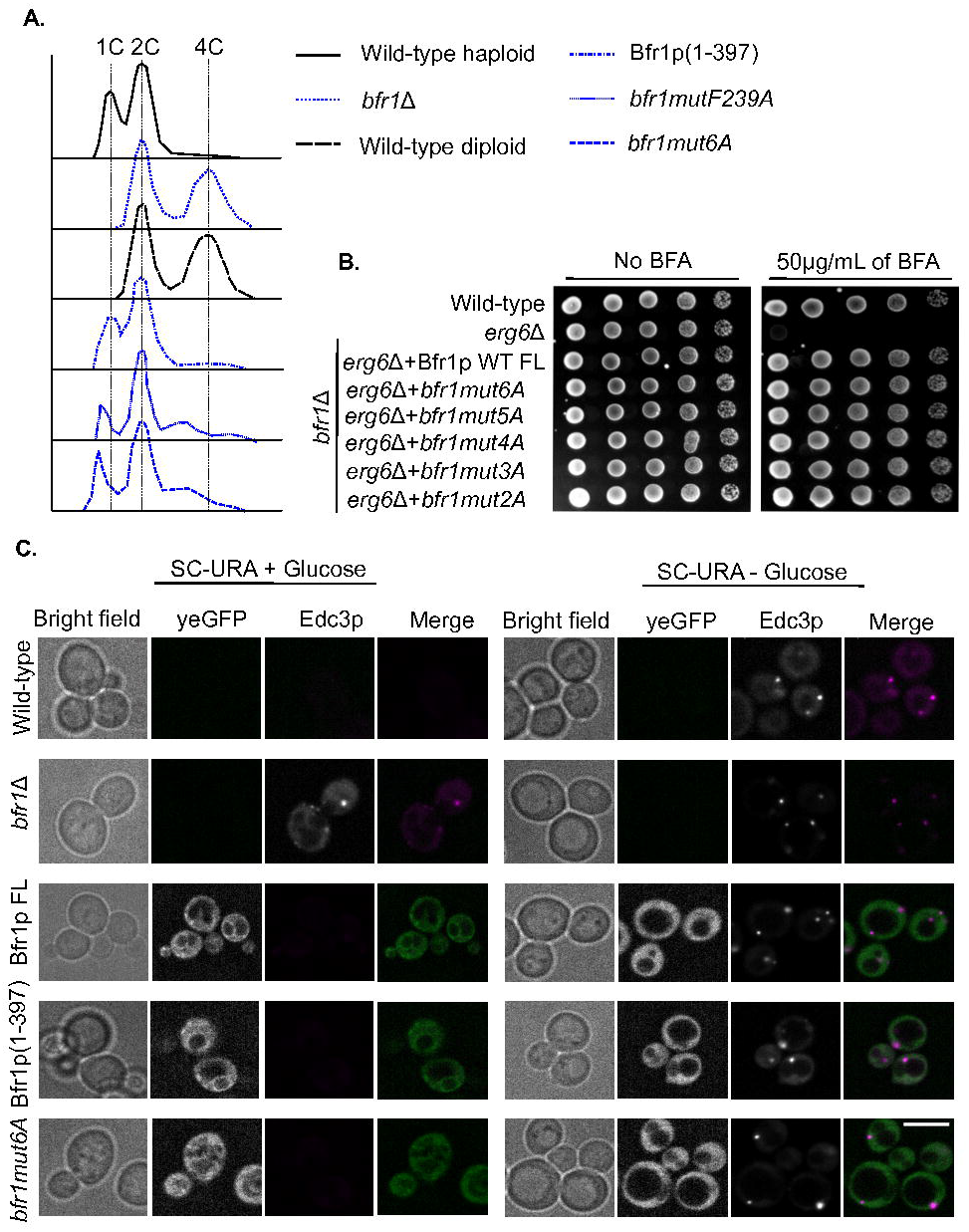
Bfr1p associated phenotypes are independent of Bfr1p-RNA interactions. A. Flow cytometry analysis of Bfr1p variants showing that ploidy is not changed by the mutations in RNA-binding residues of Bfr1p. Histograms show plots of DNA content after propidium iodide staining. B. Brefeldin A sensitivity of *erg6*Δ is rescued by overexpression of wild-type Bfr1p and RNA-binding mutants. Bfr1p mutants (*bfr1mut5A*: *R38A, K138A, F211A, Y225A, F239A*; *bfr1mut4A*: *R38A, K138A, F211A, F239A*; *bfr1mut3A*: *R38A, K138A, F239*; *bfr1mut2A*: *K138A, F239A*) were expressed from YEplac181 plasmids in cells deleted for *BFR1* and *ERG6*. Logarithmically growing cells were serially diluted, plated on agar with or without 50 µg/mL Brefeldin A and grown for 72 hrs at 30 °C. C. Premature P-bodies are not induced by RNA-binding mutations of Bfr1p. Images were collected from logarithmically growing cells shifted for 30 minutes to media with or without glucose to induce P-bodies. Edc3p-mCherry serves as P-body marker. Unlike a *BFR1* deletion, neither RNA-binding mutations nor Bfr1p (1-397) show any P-body foci in cells growing in glucose-containing medium. Scale bar, 5 µm.

Taken together, point mutations in F239 or the six amino acids that form crosslinks to RNA affect the intracellular distribution of Bfr1p but not ploidy maintenance, brefeldin A sensitivity or P-body formation, indicating that RNA-binding might not be required for these functions of Bfr1p.

### Co-localization of the Bfr1p mRNA target *ERG4* with ER is independent of Bfr1p

Several hundred mRNAs have been identified as potential binding partners of Bfr1p (Hogan et al. 2008; Mitchell et al. 2013; Lapointe et al. 2015) but it has not yet been addressed how Bfr1p affects their function. We therefore addressed the physiological changes of potential Bfr1p RNA targets upon mutation of its RNA contact sites. Since Bfr1p is enriched at the ER where mRNAs encoding membrane or secreted proteins are translated, we focused on five Bfr1p targets that encode proteins of these groups and whose functions have been linked to described phenotypes of *bfr1*Δ or Bfr1p overexpression (Setty et al. 2003; Gillingham and Munro 2003; Beh et al. 2001; Entian et al. 1999; Lai et al. 1994; Zweytick et al. 2000). These candidates were selected by comparing the top identified Bfr1p and Scp160p binders from two studies (Hogan et al. 2008; Lapointe et al. 2015). From this list we chose mRNAs that were bound by both proteins. Based on Bfr1p’s suppression of the growth inhibition by brefeldin A (Jackson and Képès 1994), the subsequent selection focused on mRNAs encoding ER- or Golgi-localized proteins (*ERG4, RUD3, SGM1*), proteins involved in ER-Golgi transport (*IMH1*) or those involved in sterol biosynthesis (*ERG4*) or binding (*OSH7*). Since loss of Bfr1p has been linked to premature P-body formation (Weidner et al. 2014), we first tested if loss of *BFR1*, deletion of the last coiled-coil domain, or mutations in RNA contact sites affect stability of the five selected mRNAs. As judged by qRT-PCR from total RNA isolates, none of the five tested mRNAs showed a significant change in their levels (Figure 3A). We verified if these mRNAs are bona-fide targets of Bfr1p and if the introduced mutations of Bfr1p RNA-binding sites interfere with their binding, we investigated two mRNAs, *OSH7* and *ERG4*. We performed co-immunoprecipitation using GFP-tagged versions of wild type Bfr1p, *bfr1mut6A*, and Bfr1p(1-397), as well as a control RNA-binding GFP fusion protein, Khd1p (Hasegawa et al. 2008; Syed et al. 2018). *MID2*, an mRNA bound by Khd1p (Syed et al. 2018) served as control mRNA. Only *ERG4* mRNA showed an enrichment in a Bfr1-GFP pulldown whereas *OSH7* was even more enriched in GFP-Khd1p pulldowns than by Bfr1-GFP pulldowns (Figure 3B). Importantly, *ERG4* enrichment is lost in the *bfrmut6A* mutant, supporting the idea that this mRNA is a direct target of Bfr1p.

**Figure 3.**
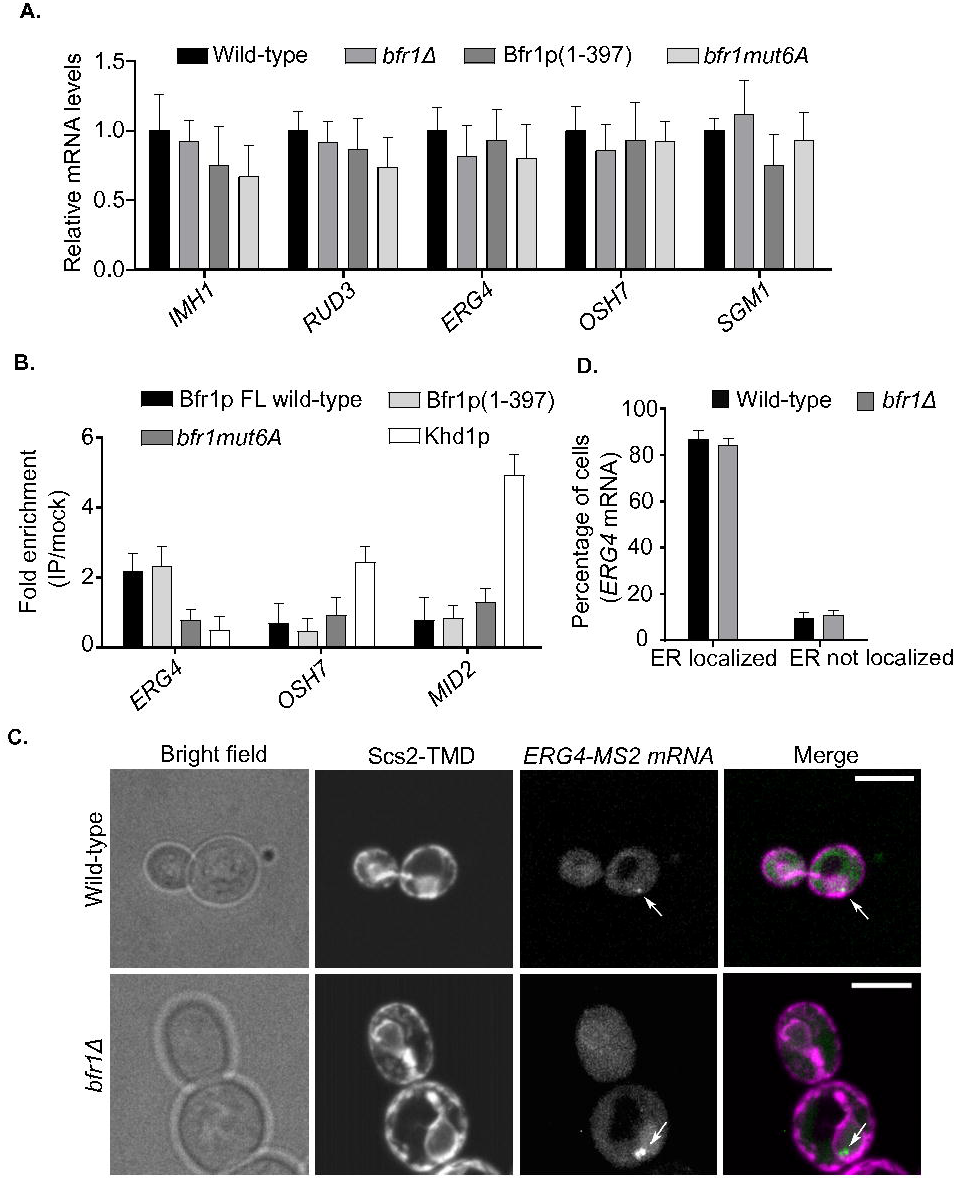
Bfr1p is not required for association of *ERG4* mRNA with ER. A. Mutations in RNA-binding sites in Bfr1p do not affect target mRNA levels. Quantification of mRNA levels of *IMH1, RUD3, ERG4, OSH7*, and *SGM1* by qRT-PCR. Data is presented as mean values from three independent experiments with ± SD. B. *ERG4* mRNA binding is strongly affected in *bfr1mut6A*. RNA-binding of full length Bfr1p (FL wild-type), Bfr1p(1-397), and *bfr1mut6A* is assessed by co-immunoprecipitation of *ERG4* and *OSH7* mRNAs and qRT-PCR. *MID2* mRNA serves as a non-target for Bfr1p. Data is presented as mean values from three biological and two technical replicates, each with ± SD. C. *ERG4* mRNA association with ER is independent of Bfr1p. Representative images of cells expressing MS2-tagged *ERG4* mRNA in wild-type and *bfr1*Δ. Scs2-TMD-2x RFP serves as ER marker. White arrows indicate *ERG4* mRNA particles. Scale bar, 8 µm. D. Quantification of *ERG4* mRNA co-localization with ER in wild-type and *bfr1*Δ. Data is presented as mean values from at least 100 cells from three biological replicates with ± SD.

We have previously shown that RBPs like She2p (Schmid et al. 2006; Fundakowski et al. 2012) or Khd1p (Syed et al. 2018) are required for ER localization of bound mRNAs. Since Bfr1p co-localizes with ER and *ERG4* encodes a membrane protein that is supposed to be synthesized from an ER-associated mRNA, we next tested if loss of *BFR1* interferes with ER association of *ERG4. ERG4* mRNA was tagged with 12 MS2 stem loops using the recently published improved MS2-tagging system (Tutucci et al. 2017) and localization of the mRNA was followed by co-expression of a GFP-MCP (MS2 coat protein) fusion protein. ER was detected by a fusion of the transmembrane domain of Scs2p (Scs2-TMD) and 2x RFP (Loewen et al. 2007). In the majority of wild-type cells (86.6+/-3.6%) and cells lacking *BFR1* (84+/-2.6 %) *ERG4-MS2* RNA particles were detected at or very close to the ER marker (Figure 3C and 3D), indicating that Bfr1p is not required for ER association of *ERG4* mRNA.

### Synthesis of the sterol reductase Erg4p depends on Bfr1p

In parallel to the RNA localization experiments, we wanted to determine if loss of *BFR1* impacts Erg4 protein localization. The C-24(28) sterol reductase Erg4p is a 7-transmembrane domain protein that catalyzes the final step of ergosterol biosynthesis in the yeast ER (Zweytick et al. 2000). Consistent with previous observations, a C-terminal fusion of Erg4p with yeast enhanced GFP (yeGFP) co-localizes with an ER marker (HDEL-DSRED; (Bevis and Glick 2002)), predominantly with perinuclear ER (Figure 4A). Although mRNA localization is unaffected, the protein level appears to be drastically reduced and no ER co-localization was detected in *bfr1*Δ cells (Figure 4A). Reduction in Erg4-yeGFP is not due to increased vacuolar protein degradation since deletion of *PEP4*, encoding a major vacuolar peptidase does not rescue this phenotype (Figure 4A). To determine, whether Erg4p is expressed at low levels or not at all, cell lysates were fractionated to enrich for membrane and cytosolic fractions. Western blot analysis shows weaker expression of Erg4-yeGFP in *bfr1*Δ cells as compared to wildtype and a shift in its distribution from the membrane to the cytosolic fraction (Figure 4C). Although *bfr1*Δ yeast share several phenotypes with *scp160*Δ cells, including increase in ploidy and premature P-body formation, deletion of *SCP160* does not affect Erg4p protein localization (Figure 4A, lower panels). This suggests that the observed low expression of Erg4p is specific to cells lacking Bfr1p.

**Figure 4.**
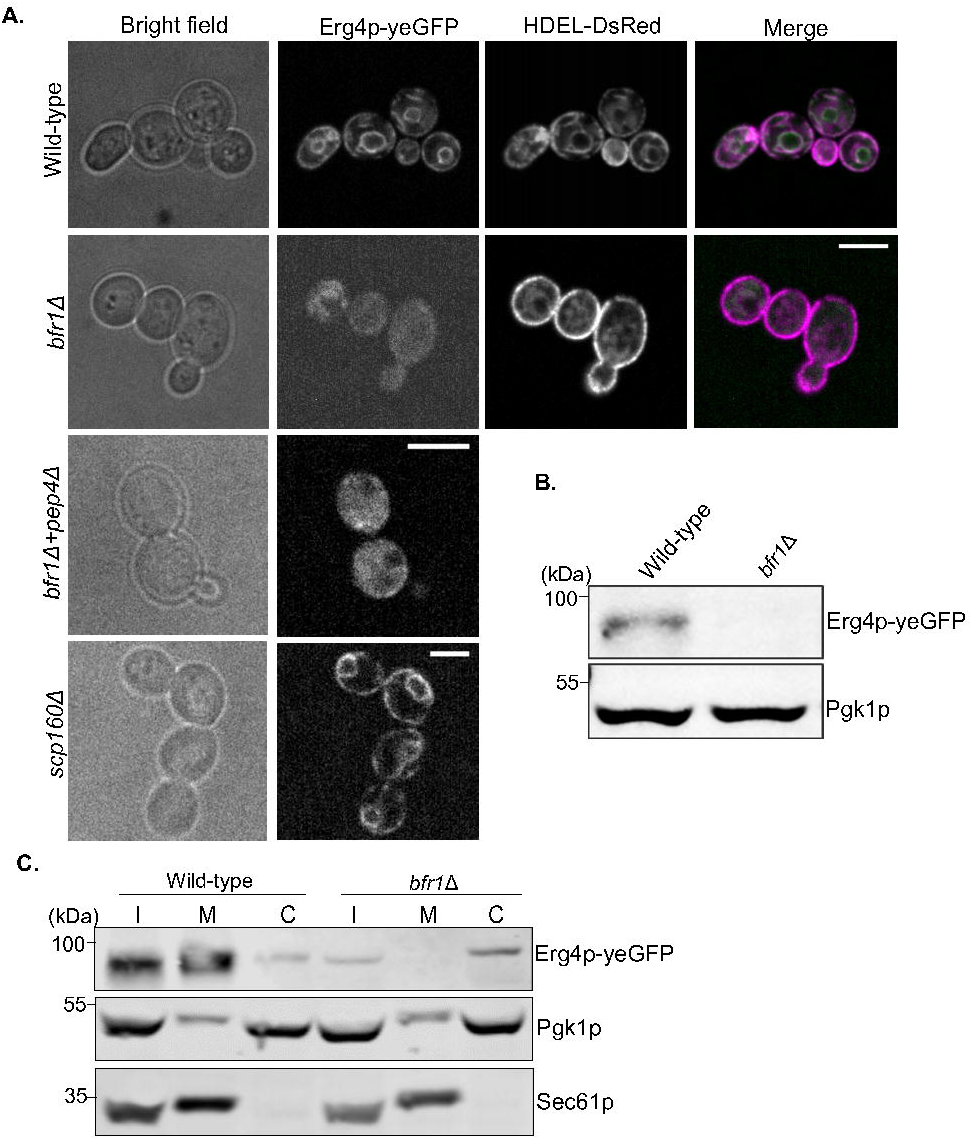
Bfr1p controls Erg4p expression and distribution. A. Erg4p distribution changes upon deletion of *BFR1*. Representative images of cells from wild-type, *bfr1*Δ, *bfr1*Δ *pep4*Δ, and *scp160*Δ strains. HDEL-DsRed serves as ER marker. Scale bar, 5 µm. B. Western-blot showing downregulation of Erg4p in *bfr1*Δ compared to wild-type. Total cell lysates were prepared from wild-type and *bfr1*Δ cells expressing a yeGFP-tagged Erg4p protein and Erg4p detected by an anti-GFP antibody. C. Erg4p shifts from ER to cytoplasm in *bfr1*Δ cells. Western blot following subcellular fractionation of wildtype and *bfr1*Δ cells expressing yeGFP-tagged Erg4p. I (input), M (membrane fraction; pellet 18000 x*g*), and C (cytoplasmic fraction; supernatant 18000 x*g*). Pgk1p and Sec61p serve as a cytoplasmic or ER marker, respectively.

### Bfr1p affects polysome association of *ERG4* mRNA

Since Bfr1p has been shown to associate with polysomes (Lang et al. 2001) we decided to test if Bfr1p directly interacts with ribosomes and if its interaction with mRNA occurs during translation. For this, we first generated a strain expressing an HA-tagged Bfr1p and showed that it co-immunoprecipitates ribosomes, using the ribosomal small subunit protein Rps3p as proxy (Figure 5A, IP). Treatment with RNAse, which depleted only mRNAs but did not affect ribosomal RNA (Figure S3A) shifts a large fraction of Rps3p from the co-immunoprecipitated fraction into the flow through (Figure 5A, FT). This indicates that to a large part binding of Bfr1p to ribosomes is mRNA dependent. To check whether Bfr1p interacts with its target mRNAs (e.g. *ERG4*) during translation or independent of it, we replaced the wild-type *ERG4* mRNA with a mutant *ERG4* lacking the AUG and expressed it in cells with a yeGFP-tagged variant of Bfr1p. Functional *ERG4* mRNA co-immunoprecipitates with Bfr1p-yeGFP (enrichment of 1.96 +/-0.34 over mock), whereas *ERG4(-AUG)* mRNA does not (1.11 +/-0.44; Figure 5B), which is consistent with the idea of translation-dependent association of *ERG4* mRNA and Bfr1p.

**Figure 5.**
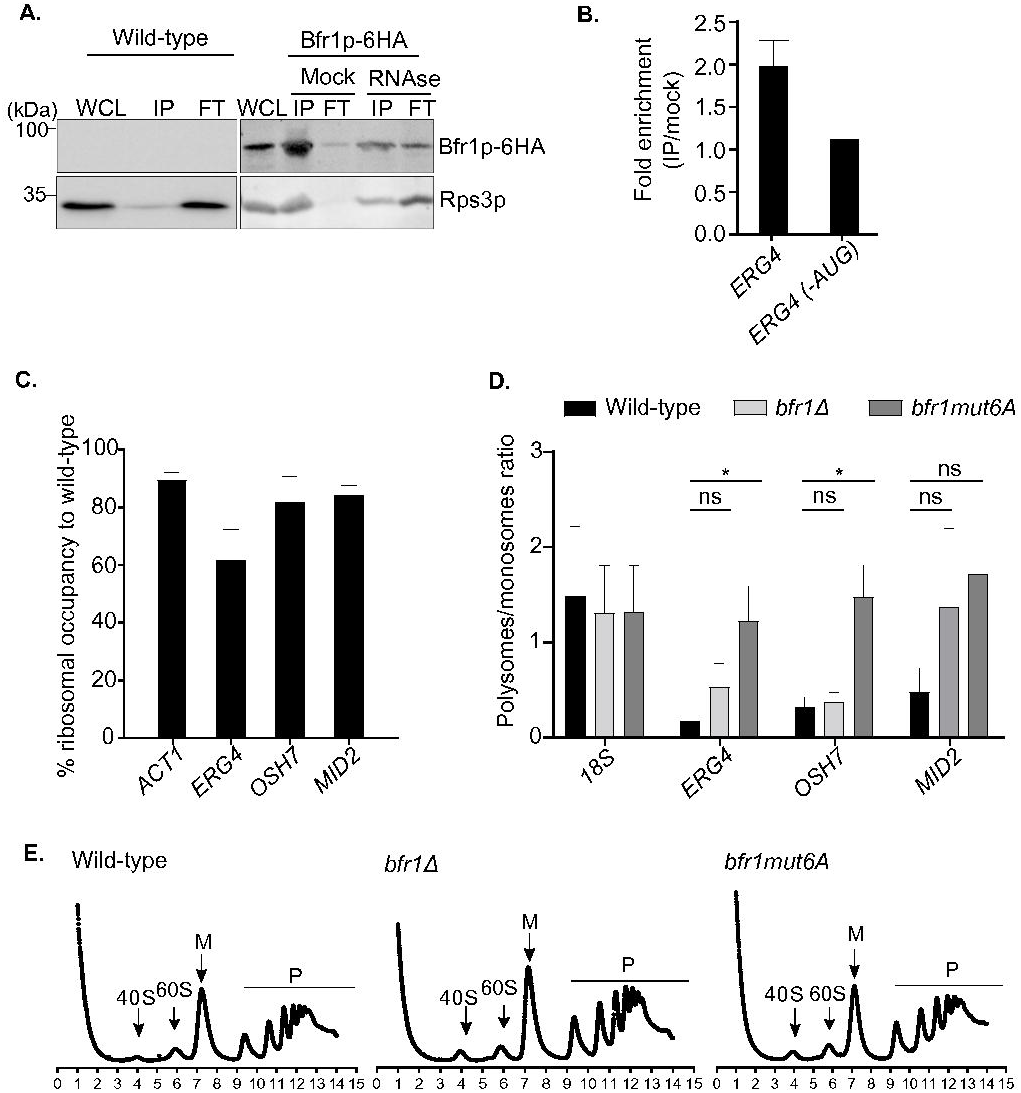
Bfr1p is required for efficient translational elongation of *ERG4* mRNA. A. Bfr1p co-precipitates with small ribosomal subunit Rps3p in and RNA dependent manner. A representative image of three independent experiments shows partial loss of Rps3p bound to Bfr1p-6HA after treatment with RNAse compared to the mock treatment. WCL: whole cell lysate, IP: immunoprecipitation, FT: flow through. B. Bfr1p interaction with *ERG4* mRNA is increased upon translation. Immunoprecipitation was performed from the lysates of cells expressing Bfr1p-yeGFP and wild-type cells to co-precipitate *ERG4* mRNA with or without AUG and levels were quantified by qRT-PCR. Binding of *ERG4* mRNA lacking an AUG is reduced by 43%. Data are represented as fold change ratio of the IP versus mock from three independent replicates with ± SD. C. Ribosomal occupancy of *ERG4* mRNA is reduced in the absence of Bfr1p. Ribosome Affinity Purification (RAP) followed by qRT-PCR was performed to measure ribosome association of *ACT1, ERG4, OSH7,* and *MID2* mRNAs in wild-type and *bfr1*Δ cells. An untagged (mock) strain was used to normalize the data. Quantification graphs show % of occupancy in *bfr1*Δ to the wild-type levels in three biological and two technical replicates of each with ± SD. D. Polysome association of *ERG4* mRNA changes in *bfr1mut6A* and *bfr1*Δ. Sucrose density gradient fractionation was used to separate monosomes and polysomes, and RNAs from the fractions were quantified by qRT-PCR. Results are displayed as the fold change ratio of polysomes/monosomes for four mRNAs (normalized to *ACT1* levels) from three independent experiments with ± SD. * indicates p <0.05 and ns = non-significant. E. Polysome profiles from *bfr1mut6A,* and *bfr1*Δ compared to wild-type cells. M (monosomes), P (polysomes).

We next investigated if loss of *BFR1* or mutations in its RNA-binding sites result in changes of translation of *ERG4*. We first addressed this by ribosomal affinity purification in combination with mRNA detection by RT-PCR (Hirschmann et al. 2014). Ribosomes and bound mRNAs were affinity-purified from cycloheximide-treated cells via a TAP-tagged ribosomal protein Rpl6p. The level of ribosome-associated *OSH7, ACT1* and *MID2* mRNAs in *bfr1*Δ cells is very similar to that of wild-type cells (Figure 5C), indicating that their ribosomal occupancy is independent of Bfr1p. In contrast, the amount of ribosome-associated *ERG4* is reduced. Since this method cannot distinguish between monosome- or polysome-associated mRNAs, we applied polysome fractionation and determined the distribution of mRNAs between mono- and polysomes, which can serve as proxy for the translational efficiency of a given mRNA. Ribosome distribution between fractions containing free ribosomal subunits, monosomes, or polysomes is similar in wildtype, *bfr1*Δ and *bfr1mut6A* cells, as judged by the polysome/monosome ratio of 18S rRNA (Figure 5D) or polysome profiles (Figure 5E). In contrast, significant changes were seen for the distribution of *ERG4* (p<0.0299) and *OSH7* (p<0.0178) between polysomes and monosomes when comparing wildtype cells and cells expressing the Bfr1p mutant *bfr1mut6A*. We also observed changes in the distribution of the control mRNA *MID2*, although with low significance (p <0.0708). Polysome profiles show *ERG4* and *OSH7* mRNAs are increased in polysomes of *bfr1mut6A* compared to wildtype cells. Although this might suggest better translation, which is in contrast to protein expression (Figure 4B). This apparent discrepancy might be due to the presence of *ERG4* and *OSH7* mRNAs on stalled ribosomes in *bfr1* mutants. A similar observation has been made in case of Scp160p where at least one of its target mRNAs, *PRY3*, is increased in polysomes in cells lacking Scp160p while Pry3 protein expression is reduced (Hirschmann et al. 2014).

## DISCUSSION

The yeast RNA-binding protein Bfr1p has been implicated in various cellular functions, ranging from control of spindle pole body duplication to P-body formation. Since Bfr1p binds to several hundred mRNAs and is found in the polysome fraction during sucrose gradient centrifugation, the multiple phenotypes associated with loss of *BFR1* might result from a function in translation or translation regulation. In order to establish additional evidence for this model and to investigate if loss of RNA-binding is causing the reported phenotypes associated with *BFR1* deletion, we studied the consequences of mutations in documented RNA-binding sites (Kramer et al. 2014). The mutated amino acids are conserved between several yeast species and are located in the first and second coiled coil domain of Bfr1p as well as in the region connecting these two coiled coils. Conversion of six of these RNA contact sites to alanines (*bfr1mut6A*) results in loss of the mRNA binding capacity as shown for *ERG4* mRNA. As a consequence, the ER localization of Bfr1p is lost and the protein redistributes to the cytoplasm. This suggest that Bfr1p is targeted to ER in a piggy-back manner via bound mRNAs that are translated at the cytoplasmic face of the ER. Among the six conserved RNA contact sites, phenylalanine 293 seems to be the most critical since loss of ER association can also be seen for the F293A mutation. Surprisingly, although ER association is lost in these mutants, none of the other described phenotypes that have been described for *bfr1*Δ cells can be detected (Jackson and Képès 1994; Weidner et al. 2014; Simpson et al. 2014). Cells expressing *bfr1mut6A* do not show loss of ploidy control and stay haploid. Like wildtype *BFR1*, overexpression of *bfr1mut6A* can fully rescue an *erg6*Δ mutant when exposed to brefeldin A (Jackson and Képès 1994). Finally, unlike *bfr1*Δ cells, cells expressing the RNA binding mutant do not show a premature formation of P-bodies in unstressed cells (Weidner et al. 2014). Thus, RNA-binding does not seem to be important for Bfr1p’s functions in ploidy control, repression of P-body formation, or brefeldin A resistance. Similarly, the third coiled coil domain of Bfr1p, which does not contain any identified RNA contact site is also dispensable for controlling P-body formation and ploidy. This suggests that these functions of Bfr1p are associated with its amino-terminal section but might be mediated by protein-protein rather than protein-RNA interactions.

A large fraction of Bfr1p target mRNAs encode proteins translated at the ER (Lapointe et al. 2015). Several RBPs have been described that participate in or mediate mRNA association with the ER independent of translation or SRP-mediated targeting (Cui and Palazzo 2014). These include mammalian p180 and Kinectin (CUI 2012) and the yeast RBPs She2p (Schmid et al. 2006) or Khd1p (Syed et al. 2018). In contrast to these, Bfr1p is not required for ER localization of at least one of its bound mRNAs, *ERG4*. In this respect, it mimics Scp160p, with which it shares its polysome association and ER co-localization. Although not necessary for mRNA localization, Bfr1p is required for the proper translation of the encoded Erg4 protein, which is consistent with its binding to translating polyribosomes ((Lang et al. 2001); Fig. 5A+B). Not only is the total amount of Erg4 protein lower in *bfr1*Δ cells but the distribution of this membrane protein is altered and enriched in the cytosol rather than the membrane fraction. Erg4p catalysis the final step of ergosterol synthesis from 5,7,22,24(28)-ergostatetraenol to ergosta-5,7,22-trien-3beta-ol. However, despite the observed altered protein distribution, we have not detected changes in the ergosterol level of *bfr1*Δ cells versus wild-type cells. It is thus likely that the remaining amounts of properly translated and targeted Erg4p suffices for ergosterol synthesis.

Concomitant with a postulated role of Bfr1p in translation of *ERG4* mRNA, we also observed a decrease of the fraction of *ERG4* mRNA that co-purifies with ribosomes. Surprisingly, the ratio of *ERG4* mRNA on polysomes versus monosomes increases, which is apparently contradictory to the findings of reduced translation. However, a similar observation has been made for cells lacking the conserved RBP Scp160p, although for a different set of mRNAs (Hirschmann et al. 2014). The reduced association of certain tRNAs with translating ribosomes in cells lacking Scp160p has led to the model that Scp160p is required for efficient translation of specific mRNA subsets and that its loss results in pausing or stalling of ribosomes on mRNAs (Hirschmann et al. 2014). We envision that Bfr1p might function similarly to Scp160p but might be important for a regulating a different set of mRNAs encoding membrane or secreted proteins at ER.

## MATERIALS AND METHODS

### Yeast strains and plasmids

Generation of yeast strain and plasmids used in this study as well as general methods including cell lysis and subcellular fractionation are described in Supplementary Methods.

### Confocal microscopy

*In vivo* imaging of fluorescent labelled proteins was carried out from inoculation of single colonies into SC or SDC medium containing 2% glucose and overnight growth at 30° C. Logarithmically growing cells were harvested and re-suspended in 100µl of fresh medium. Cells were spread on thin agarose pads containing SC or SDC with 2% glucose and grown for 30 minutes at 30° C before observing in microscope (ZEISS AxioExaminer equipped with a CSU spinning disc confocal unit; Visitron Systems, Germany). Images were acquired with a 63x oil objective using VisiView software (Visitron Systems). Post image processing was performed using Fiji software as described in (Syed et al. 2018; Hermesh et al. 2014). For *ERG4* mRNA localization, images were acquired, with each containing at least 60 cells. For P-bodies analysis, logarithmically growing cells were harvested, washed once with sterile water before cells were spread on agarose pads containing SDC medium without glucose and grown at 30°C for 30 min to induce P-bodies.

### Co-immunoprecipitations

For protein-mRNA co-precipitations, experiments were performed as described (Syed et al. 2018) with the following changes. 100 OD_600_ units of cells were harvested and resuspended in 8-10 µl/OD_600_ of lysis buffer (10 mM Tris-HCl pH 7.5, 150 mM NaCl, 2 mM EDTA, 0.1% Triton X-100, 1x Protease inhibitor cocktail and 0.5 U/µL Ribolock RNase inhibitor). Glass bead lysis was performed and cell debris was removed by 5 minutes centrifugation at 5000 *x*g at 4° C. Protein concentrations were measured and 200 µg of lysates were subjected to immunoprecipitation with GFP-Trap®_MA magnetic beads (Chromotek) at 4° C for two hours on a rotating wheel. The beads were blocked prior to the immunoprecipitations with blocking buffer (10 mM Tris-HCl pH 7.5, 150 mM NaCl, 2 mM EDTA, 0.1% Triton X-100, 0.1 mg/ml *E.coli* tRNA and 0.4 mg/ml Heparin). Captured beads were washed 3x with 500 µl of wash buffer (10 mM Tris-HCl pH 7.5, 150 mM NaCl, 2 mM EDTA, and 0.1% Triton X-100) and resuspended beads with 125 µl of HPLC grade water (Sigma). 75µl of the bead slurry was used for SDS-PAGE and western blotting. RNA extraction was carried out with the remaining 50 µl of beads, to which 50 ng of spike RNA (*in vitro* transcribed *Arabidopsis* phosphoribulokinase RNA) had been added. For extraction 5 µl of 3 M Sodium acetate pH 5.2, 2.5 µl of 20% SDS and 50 µl of phenol/chloroform/isoamylalcohol (PCI) pH 4.5 (Roth, Germany) was added and the mixed sample centrifuged at maximum speed for 30 min at 4° C. Nucleic acids were precipitated overnight at −80° C with 20 µg of glycogen and 100 µl of 96% ethanol and RNA was resuspended in 20 µl of RNAse free water before continuing with cDNA synthesis and qPCR.

Co-immunoprecipitations of Rps3p with Bfr1p-3HA were performed with similar conditions to Bfr1p-yeGFP but immunoprecipitated with Protein G Sepharose^®^ 4 Fast Flow beads (GE Healthcare). Prior to the immunoprecipitations, protein concentration was determined and 125 µg of lysates were prepared. RNAse treatment was performed with 100 µg/ml of RNAse A for 30 min at 37° C. Lysates were pre-cleared with 20 µl of Protein G Sepharose beads overnight 4° C on a rotating wheel. Beads were removed by centrifugation (twice at 1,000 x*g*) and then pre-cleared lysates were supplemented with anti-HA high affinity monoclonal antibody (clone 3F10, Roche) at 2 µg/ml final concentration along with 20 µl Protein G Sepharose beads and incubated at 4° C for 4 hours on a rotating wheel. After washing, beads were resuspended in 1x Laemmli buffer and continued with SDS-PAGE and western blot analysis.

### RNA extraction, RT-PCR and qRT-PCR

Total RNA was extracted from 20 OD_600_ units of yeast strains RJY358 and RJY4626. Glass beads lysis was performed in 500 µl of Cross buffer I (0.3 M NaCl, 10 mM Tris-what pH 7.5, 1 mM EDTA, 0.2 % SDS) and 400 µl of phenol:chloroform:isoamylalcohol by four pulses of 120 seconds with breaks of 60 seconds on ice. After centrifugation for 30 minutes at full speed the upper phase was transferred to a new tube and nucleic acids precipitated with ethanol for 30 min at −20° C. Total RNA pellets were washed and resuspended in 20 µl of HPLC grade water before proceeding to cDNA synthesis and qPCR.

Reverse transcription reactions for all the experiments were performed as described in (Syed et al. 2018) with some modifications. In brief, 1 µg of RNA samples were treated with RQ1 DNase (Promega) and reverse transcription reactions performed using the High-Capacity cDNA Reverse Transcription Kit (Applied Biosystems). Quantitative RT-PCR (qRT-PCR) was performed in 10 µl reactions containing 2.5 µl of cDNA samples at 1:25 or 1:50 dilutions. Target specific primers were designed with Primer3 software (Rozen, Skaletsky 2000). All reactions were performed from a minimum of three biological replicates and two technical replicates of each. Quantification was performed by comparative ΔΔCt method (Livak and Schmittgen 2001; Syed et al. 2018) and values were normalized via the spiked-in RNA or via *ACT1* mRNA signals. For the RT-PCR reactions after RNAse treatment, a PCR reaction with 1 µl of cDNA with dilutions up to 1:1000 was performed using Taq DNA polymerase (Genaxxon Bioscience). PCR was performed for 18 cycles using primers specific to *18S* rRNA and *ERG4* mRNA and amplified products were separated by 1% agarose gel electrophoresis.

### Ribosome affinity purifications (RAP-IP)

Ribosome affinity purification (RAP-IP) to determine *ERG4* and *OSH7* mRNAs bound to ribosomes was performed as described in (Hirschmann et al. 2014). For the quantifications of mRNAs bound to the ribosomes, qRT-PCR was performed and the enrichment of mRNAs determined using the ΔΔCT method. An enrichment of mRNAs from TAP purification (in wild-type and *bfr1*Δ) was considered only if at least two fold greater than from mock purification (strains without TAP tags). The % ribosomal occupancy of *bfr1*Δ was then plotted against wild-type levels for the *ACT1, ERG4, MID2* and *OSH7* mRNAs (Fig 5C).

### Brefeldin A sensitivity drop assay

A single colony of yeast cells was inoculated and grown overnight before diluting it into fresh medium to grow until logarithmic phase. Cells were harvested and washed once with sterile water. One OD_600_ unit of cells was used for serial dilution and 3 µl of dilutions were plated on SDC-leu medium with or without 50 µg/ml of brefeldin A (eBioscience^TM^). Cells were grown for 72 hrs at 30°C.

### Ploidy analysis by flow cytometry

Cells were prepared for flow cytometry analysis as described (Baum et al. 2004; Hirschmann et al. 2014). Propidium iodide fluorescence was detected in a Beckman Coulter CytoFlex LX system with a 675/30 nm filter. Data was analyzed using CytExpert software (Beckman Coulter) and representative graphs were plotted manually using values obtained from the software.

### Polysomes profiling

Separation of mono- and polysomes was done as described in (Mittal et al. 2017) and performed with three biological replicates of wildtype, *bfr1*Δ and *bfr1mut6A* strains. Logarithmically growing cells were treated with 100µg/ml cycloheximide (CHX) for 1 minute, harvested by vacuum filtration and flash frozen with liquid nitrogen. Cells lysis was performed under cryogenic conditions using lysis buffer (20 mM Tris-HCl, pH 7.5, 100 mM NaCl, 10 mM MgCl_2_, 1% Triton X-100, 0.5 mM DDT and 100 µg/ml of CHX) and a bead mill (Spex Inc., Metuchen, NJ, USA). Cell debris was removed by centrifugation for 3 min at 3,000 *x*g and 4 °C followed by 10,000 *x*g for 5 minutes at 4 °C. To separate monosomes and polysomes, 10 A_260_ units of lysates were loaded on pre-cooled 12 ml of 7% - 47% linear sucrose gradients (50 mM Tris-HCl, pH 7.5, 50 mM NH_4_Cl, 12 mM MgCl_2,_ 0.5 mM DTT, and 100 µg/ml CHX) and centrifuged at 35,000 rpm for 3 hours at 4° C in a TH-641 rotor (Thermo Scientific) before collecting the monosome and polysome peaks (Fig 5E). RNA was isolated by adding 5µl of glycogen and phenol:chloroform (5:1) before re-extraction of the aqueous phase with chloroform and precipitation in ethanol overnight at −20°C. RNA pellets were resuspended in 30 µl of HPLC grade water and processed for cDNA synthesis and qRT-PCR.

## ACKNOWLEDGEMENTS

We are thankful to Robert H. Singer and Roy Parker for providing plasmids MCPNLSSV40-2x-yeGFP and Edc3-mCherry, respectively. We are grateful to Syed Muhammad Ibrahim for helping with subcellular fractionation, microscopy experiments and for providing spike RNA. Joyita Mukherjee helped in data analysis of qRT-PCR and co-immunoprecipitation experiments. Mathew Cheng is thanked for suggestions on the manuscript.

## REFERENCES

Auweter SD, Oberstrass FC, Allain FHT. 2006. Sequence-specific binding of single-stranded RNA: is there a code for recognition? Nucleic Acids Research 34: 4943–4959.

Baltz AG, Munschauer M, Schwanhäusser B, Vasile A, Murakawa Y, Schueler M, Youngs N, Penfold-Brown D, Drew K, Milek M, et al. 2012. The mRNA-bound proteome and its global occupancy profile on protein-coding transcripts. Molecular Cell 46: 674–690.

Baum S, Bittins M, Frey S, Seedorf M. 2004. Asc1p, a WD40-domain containing adaptor protein, is required for the interaction of the RNA-binding protein Scp160p with polysomes. Biochem J 380: 823–830.

Beckmann BM, Horos R, Fischer B, Castello A, Eichelbaum K, Alleaume A-M, Schwarzl T, Curk T, Foehr S, Huber W, et al. 2015. The RNA-binding proteomes from yeast to man harbour conserved enigmRBPs. Nat Commun 6: 10127.

Beh CT, Cool L, Phillips J, Rine J. 2001. Overlapping functions of the yeast oxysterol-binding protein homologues. Genetics 157: 1117–1140.

Bevis BJ, Glick BS. 2002. Rapidly maturing variants of the Discosoma red fluorescent protein (DsRed). Nat Biotechnol 20: 83–87.

Björk P, Wieslander L. 2017. Integration of mRNP formation and export. Cell Mol Life Sci 74: 2875–2897.

Castello A, Fischer B, Eichelbaum K, Horos R, Beckmann BM, Strein C, Davey NE, Humphreys DT, Preiss T, Steinmetz LM, et al. 2012. Insights into RNA biology from an atlas of mammalian mRNA-binding proteins. Cell 149: 1393–1406.

Castello A, Fischer B, Frese CK, Horos R, Alleaume A-M, Foehr S, Curk T, Krijgsveld J, Hentze MW. 2016. Comprehensive Identification of RNA-Binding Domains in Human Cells. Molecular Cell 63: 696–710.

Castello A, Hentze MW, Preiss T. 2015. Metabolic Enzymes Enjoying New Partnerships as RNA-Binding Proteins. Trends Endocrinol Metab 26: 746–757.

Cui XA, Palazzo AF. 2014. Localization of mRNAs to the endoplasmic reticulum. WIREs RNA 5: 481–492.

Dreyfuss G, Kim VN, Kataoka N. 2002. MESSENGER-RNA-BINDING PROTEINS AND THE MESSAGES THEY CARRY. Nature Reviews Molecular Cell Biology 3: 195–205.

Entian KD, Schuster T, Hegemann JH, Becher D, Feldmann H, Guldener U, Götz R, Hansen M, Hollenberg CP, Jansen G, et al. 1999. Functional analysis of 150 deletion mutants in Saccharomyces cerevisiae by a systematic approach. Mol Gen Genet 262: 683–702.

Fundakowski J, Hermesh O, Jansen R-P. 2012. Localization of a subset of yeast mRNAs depends on inheritance of endoplasmic reticulum. Traffic 13: 1642–1652.

Gillingham AK, Munro S. 2003. Long coiled-coil proteins and membrane traffic. Biochim Biophys Acta 1641: 71–85.

Graham TR, Scott PA, Emr SD. 1993. Brefeldin A reversibly blocks early but not late protein transport steps in the yeast secretory pathway. EMBO J 12: 869–877.

Hasegawa Y, Irie K, Gerber AP. 2008. Distinct roles for Khd1p in the localization and expression of bud-localized mRNAs in yeast. RNA 14: 2333–2347.

Hentze MW, Castello A, Schwarzl T, Preiss T. 2018. A brave new world of RNA-binding proteins. Nature Reviews Molecular Cell Biology 546: 617.

Hermesh O, Genz C, Yofe I, Sinzel M, Rapaport D, Schuldiner M, Jansen RP. 2014. Yeast phospholipid biosynthesis is linked to mRNA localization. Journal of Cell Science 127: 3373–3381.

Hirschmann WD, Westendorf H, Mayer A, Cannarozzi G, Cramer P, Jansen RP. 2014. Scp160p is required for translational efficiency of codon-optimized mRNAs in yeast. Nucleic Acids Research 42: 4043–4055.

Hogan DJ, Riordan DP, Gerber AP, Herschlag D, Brown PO. 2008. Diverse RNA-Binding Proteins Interact with Functionally Related Sets of RNAs, Suggesting an Extensive Regulatory System. PLoS Biol 6: e255.

Jackson CL, Képès F. 1994. BFR1, a multicopy suppressor of brefeldin A-induced lethality, is implicated in secretion and nuclear segregation in Saccharomyces cerevisiae. Genetics 137: 423–437.

Jan CH, Williams CC, Weissman JS. 2014. Principles of ER cotranslational translocation revealed by proximity-specific ribosome profiling. Science 346: 1257521.

Kramer K, Sachsenberg T, Beckmann BM, Qamar S, Boon K-L, Hentze MW, Kohlbacher O, Urlaub H. 2014. Photo-cross-linking and high-resolution mass spectrometry for assignment of RNA-binding sites in RNA-binding proteins. Nature Methods 11: 1064–1070.

Kwon SC, Yi H, Eichelbaum K, Föhr S, Fischer B, You KT, Castello A, Krijgsveld J, Hentze MW, Kim VN. 2013. The RNA-binding protein repertoire of embryonic stem cells. Nature Structural & Molecular Biology 20: 1122–1130.

Lai MH, Bard M, Pierson CA, Alexander JF, Goebl M, Carter GT, Kirsch DR. 1994. The identification of a gene family in the Saccharomyces cerevisiae ergosterol biosynthesis pathway. Gene 140: 41–49.

Lang BD, Fridovich-Keil JL. 2000. Scp160p, a multiple KH-domain protein, is a component of mRNP complexes in yeast. Nucleic Acids Research 28: 1576–1584.

Lang BD, Li Am, Black-Brewster HD, Fridovich-Keil JL. 2001. The brefeldin A resistance protein Bfr1p is a component of polyribosome-associated mRNP complexes in yeast. Nucleic Acids Research 29: 2567–2574.

Lapointe CP, Wilinski D, Saunders HAJ, Wickens M. 2015. Protein-RNA networks revealed through covalent RNA marks. Nature Publishing Group 12: 1163–1170.

Lewis HA, Musunuru K, Jensen KB, Edo C, Chen H, Darnell RB, Burley SK. 2000. Sequence-specific RNA binding by a Nova KH domain: implications for paraneoplastic disease and the fragile X syndrome. Cell 100: 323–332.

Livak KJ, Schmittgen TD. 2001. Analysis of Relative Gene Expression Data Using Real-Time Quantitative PCR and the 2-ΔΔCT Method. Methods 25: 402–408.

Loewen CJR, Young BP, Tavassoli S, Levine TP. 2007. Inheritance of cortical ER in yeast is required for normal septin organization. The Journal of Cell Biology 179: 467–483.

Low YS, Bircham PW, Maass DR, Atkinson PH. 2014. Kinetochore genes are required to fully activate secretory pathway expansion in S. cerevisiae under induced ER stress. Mol BioSyst 10: 1790–1802.

Lu D, Searles MA, Klug A. 2003. Crystal structure of a zinc-finger-RNA complex reveals two modes of molecular recognition. Nature 426: 96–100.

Lunde BM, Moore C, Varani G. 2007. RNA-binding proteins: modular design for efficient function. Nature Reviews Molecular Cell Biology 8: 479–490.

Mitchell SF, Jain S, She M, Parker R. 2013. Global analysis of yeast mRNPs. Nature Structural & Molecular Biology 20: 127–133.

Mittal N, Guimaraes JC, Gross T, Schmidt A, Vina-Vilaseca A, Nedialkova DD, Aeschimann F, Leidel SA, Spang A, Zavolan M. 2017. The Gcn4 transcription factor reduces protein synthesis capacity and extends yeast lifespan. Nat Commun 8: 457.

Neriec N, Percipalle P. 2018. Sorting mRNA Molecules for Cytoplasmic Transport and Localization. Front Genet 9: 510.

Oubridge C, Ito N, Evans PR, Teo CH, Nagai K. 1994. Crystal structure at 1.92 A resolution of the RNA-binding domain of the U1A spliceosomal protein complexed with an RNA hairpin. Nature 372: 432–438.

Ryter JM, Schultz SC. 1998. Molecular basis of double-stranded RNA-protein interactions: structure of a dsRNA-binding domain complexed with dsRNA. EMBO J 17: 7505–7513.

Schmid M, Jaedicke A, Du T-G, Jansen R-P. 2006. Coordination of endoplasmic reticulum and mRNA localization to the yeast bud. CURBIO 16: 1538–1543.

Setty SRG, Shin ME, Yoshino A, Marks MS, Burd CG. 2003. Golgi recruitment of GRIP domain proteins by Arf-like GTPase 1 is regulated by Arf-like GTPase 3. CURBIO 13: 401–404.

Sezen B, Seedorf M, Schiebel E. 2009. The SESA network links duplication of the yeast centrosome with the protein translation machinery. Genes & Development 23: 1559–1570.

Shah N, Klausner RD. 1993. Brefeldin A reversibly inhibits secretion in Saccharomyces cerevisiae. Journal of Biological Chemistry 268: 5345–5348.

Simpson CE, Lui J, Kershaw CJ, Sims PFG, Ashe MP. 2014. mRNA localization to Pbodies in yeast is biphasic with many mRNAs captured in a late Bfr1pdependent wave. Journal of Cell Science 127: 1254–1262.

Syed MI, Moorthy BT, Jenner A, Fetka I, Jansen R-P. 2018. Signal sequence-independent targeting of MID2 mRNA to the endoplasmic reticulum by the yeast RNA-binding protein Khd1p. FEBS Letters 592: 1870–1881.

Tutucci E, Vera M, Biswas J, Garcia J, Parker R, Singer RH. 2017. An improved MS2 system for accurate reporting of the mRNA life cycle. Nature Methods 9: 777.

Weidner J, Wang C, Prescianotto-Baschong C, Estrada AF, Spang A. 2014. The polysome-associated proteins Scp160 and Bfr1 prevent P body formation under normal growth conditions. Journal of Cell Science 127: 1992–2004.

Xue Z, Shan X, Sinelnikov A, Mélèse T. 1996. Yeast mutants that produce a novel type of ascus containing asci instead of spores. Genetics 144: 979–989.

Zweytick D, Hrastnik C, Kohlwein SD, Daum G. 2000. Biochemical characterization and subcellular localization of the sterol C-24(28) reductase, erg4p, from the yeast saccharomyces cerevisiae. FEBS Letters 470: 83–87.

