## Supplemental Figures and Tables for "Local translation of yeast *ERG4* mRNA at the endoplasmic reticulum requires the brefeldin A resistance protein Bfr1"

**Supplementary Information (Manchalu et al., 2019):**

**Supplementary Figures** pages 2 - 4

**Supplementary Methods** pages 5 - 7

**Supplementary Tables** pages 8 - 12

**Figure S1.**

**A.**

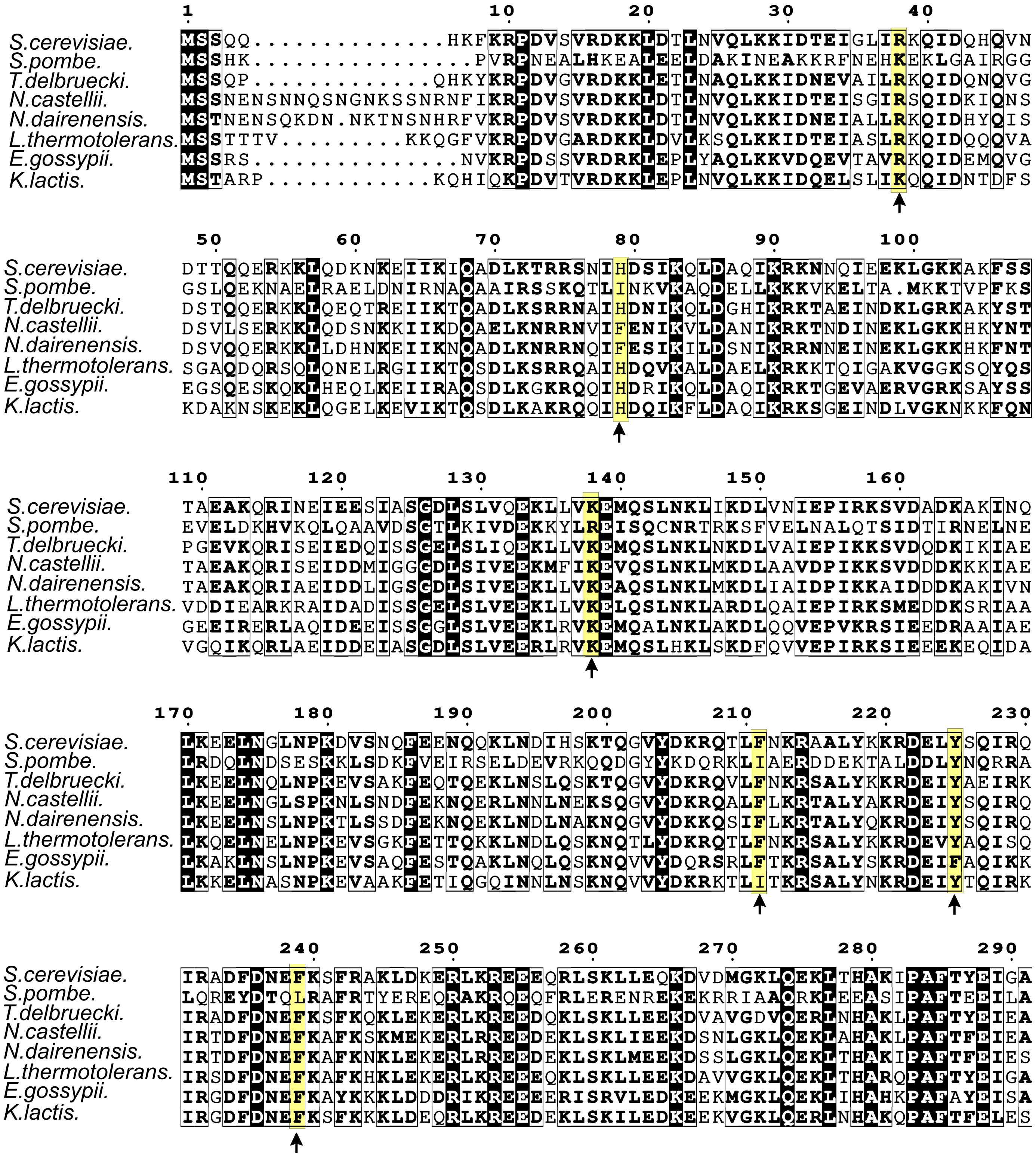


**Figure S1. Multiple-sequence alignment of Bfr1p from various fungi reveals K138 and F239 RNA-binding residues are highly conserved.**

**A.** Alignment of Bfr1p was performed for 8 different species of the fungal kingdom with Clustal Omega (EMBL-EBI). The output file was prepared with ESprint 3 and manually edited for residues 1-291 of Bfr1p. UV cross-linked RNA-binding residues of Bfr1p are highlighted in yellow with balck arrows.

**Figure S2.**

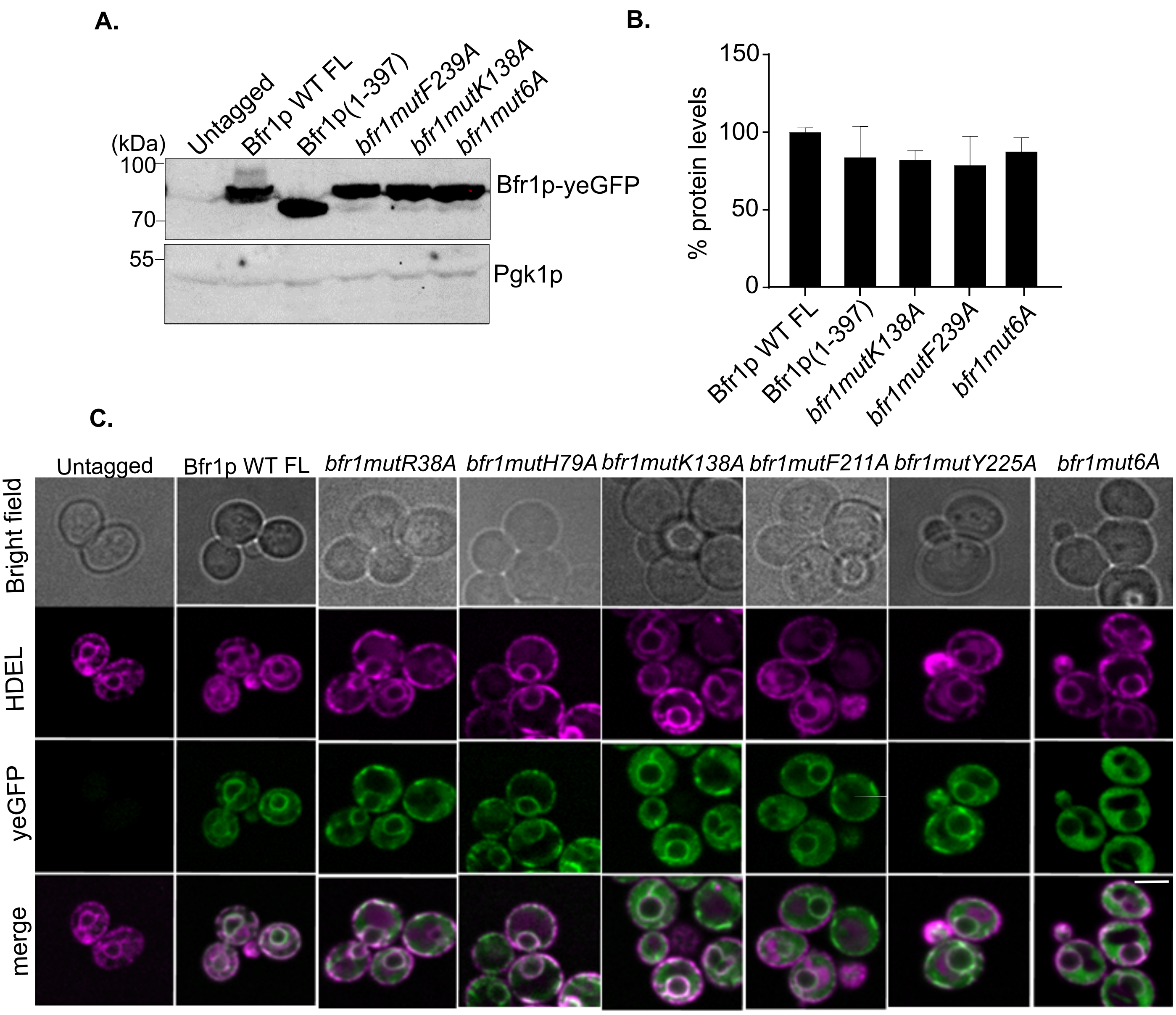


**Figure S2. Expression and localization of Bfr1p in cells with mutations in individual Bfr1p RNA-binding residues.**

**A.** Equal expression of Bfr1p in various mutants of RNA-binding residues. Total cell lysates from the yeGFP tagged Bfr1p full length (WT FL), Bfr1p (1-397), *bfr1mutF239A*, *bfr1mutK138A*, and *bfr1mut6A* were analyzed by western blotting. Pgk1p serves as a loading control and and untagged wild-type strain serves as specificity control of anti-GFP antibody.

**B.** Quantification of Bfr1p protein amounts in the western blots. Data displayed as % protein levels compared to the wild-type from three biological replicates. Normalization to Pgk1p levels was done for each strain before normalization to wildtype levels were made. Error bars with ± SD.

**C.** Bfr1p-ER co-localization images from cells expressing individual mutations in RNA-binding residues of Bfr1p with C-terminal yeGFP. HDEL-DsRed serves as ER marker. Cells with displayed single mutations do not affect ER localization of Bfr1p compared to cells expressing all 6 mutations and mutation F239A (Figure 1). Scale bar, 8 µm.

**Figure S3.**

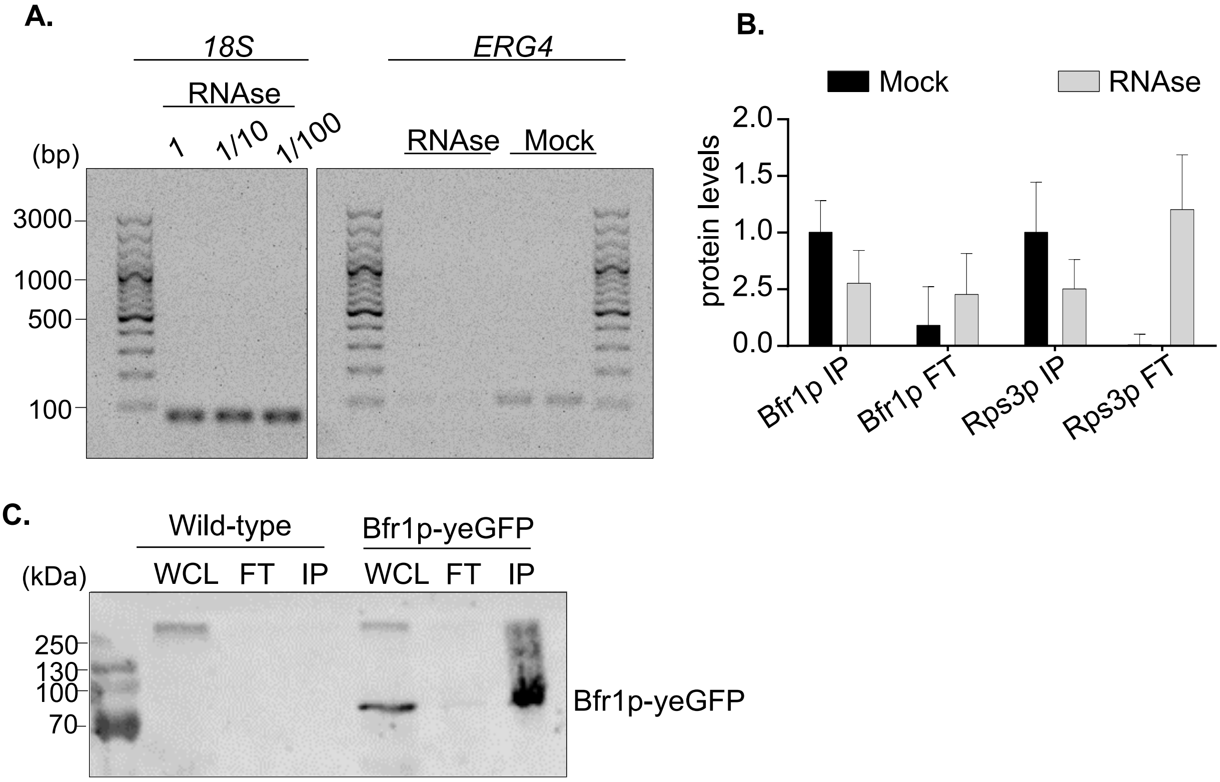


**Figure S3. Supplementary figure related to figure 5.**

**A.** Figure related to main figure 5A, showing RT-PCR amplifications of *18S* rRNA and *ERG4* mRNA treated with or without RNAse of 100µg/mL for 30 minutes. Treatment does not affect *18S* rRNA, indicating that ribosomes are still intact while mRNAs are degraded.

**B.** Western blot quantifications of immunoprecipitated Bfr1p-6HA and Rps3p from figure 5A. Data are presented for detected Bfr1p and Rps3p in IP and FT fractions that were treated with or without RNAse. Shown are mean values from three independent experiments with ± SD. The increase of Rps3p in the FT fraction after RNAse treatment indicates that Bfr1p and Rps3p interacts in RNA dependent manner. IP: immunoprecipitation, FT : flow through.

**C.**Western blot of immunoprecipitated Bfr1p-yeGFP related to main figure 5B. Wildtype (untagged) strain was used as a negative control for the immunoprecipitations. WCL: whole cell lysates, IP: immunoprecipitation, FT: flow through.

### Supplementary methods

#### Yeast strain construction

All yeast strains are derived from *W303a* background and are listed in supplementary table ST1. Gene deletion or gene tagging was performed as described (Janke et al. 2004; Gietz and Schiestl 2007) using standard PCR amplifications of desired products followed by transformation into yeast cells. The strain with BFR1 point mutations was generated as follows. We first mutated F239A using the genomic DNA from strain RJY4680 (Bfr1p-yeGFP::*HIS3MX6*) as a template and oligonucleotide pairs RJY4918 and RJY5248 (from 5'UTR region of BFR1 to F239 position), and RJY5249 and RJY3008 (from F239 position to 3'UTR region of BFR1). These two separate PCR products were then amplified in an overlap extension PCR reaction using the oligonucleotides RJY4918 and RJY3008. This intermediate PCR product served as a template for the subsequent introduction of individual mutations (R38A, H79A, K138A, F211A) using mutagenic oligonucleotides (see supplementary table ST3). After generating the final product with all six mutations the linear DNA was transformed into RJY358 and the strain RJY5204 *(bfr1mut6A*-yeGFP*::HIS3MX6*) was generated.

Strains RJY5260 (*ERG4-12xMS2V6::KanMx4*) and RJY5262 (*bfr1Δ*::*HIS3MX6*, *ERG4-12xMS2V6::KanMx4)* were generated as described (Haim-Vilmovsky and Gerst 2009; Tutucci et al. 2017). *BFR1* gene deletion strains used to express Bfr1p from plasmids were created after transformation of the corresponding plasmids to avoid diploidization as often seen in *bfr1*Δ.

#### Plasmid construction

All plasmids from this study are listed in supplementary table ST2. The oligonucleotides that were used for cloning, gene deletion, gene tagging, and introduction of single mutations are listed in supplementary table S3. Correct construction of plasmids was confirmed by restriction digestion and sequencing. The following plasmids were obtained from colleagues (RJP1870: Maya Schuldiner, RJP2121: Robert Singer, RJP1809: Roy Parker, RJP1686: Tim Levine). Plasmid RJP2000 (pRS316-pBFR1-Bfr1 FL-yeGFP) was created as follows. First, a PCR product ranging from the *BFR1* promoter region to the C-terminal yeGFP was generated using oligonucleotides RJO6367 and RJO5432 and genomic DNA of RJY4680 (Bfr1p-yeGFP::*HIS3MX6*) as a template. The product was digested with restriction enzymes *Kpn*I and *Nhe*I whose sites were introduced by oligonucleotides. RJP148 (pRS316) was digested with *Kpn*I and *Xba*I and both fragments ligated. The 3'UTR region of *BFR1* was amplified separately with oligonucleotides RJO5587 and RJO5588 using genomic DNA of strain RJY358 and digested with restriction enzymes *Eag*I and *Sac*I whose restriction sites were introduced with the oligonucleotides. The intermediate construct was digested with restriction enzymes *Eag*I and *Sac*I and the digested PCR product was ligated in to generate RJP2000.

Plasmids containing the complete *BFR1* gene with point mutations (RJP2199; *bfr1mutR38A*-yeGFP, RJP2200; *bfr1mutH79A*-yeGFP, RJP2061; *bfr1mutK138A*-yeGFP, RJP2201; *bfr1mutF211A*-yeGFP, RJP2202; *bfr1mutY225A*-yeGFP and RJP2062; *bfr1mutF239A*-yeGFP) were created from a pRS316 vector carrying *BFR1* by a PCR based site-directed mutagenesis with respective oligonucleotides.

High copy number plasmids carrying *BFR1* or its mutant constructs were created as follows. Firstly, full length BFR1 fused to the yeGFP coding region was released from RJP2000 (pRS316-pBFR1-Bfr1 FL-yeGFP) by digestion with *Kpn*I and *Sac*I and ligated into the vector YEplac181 to create plasmid YEplac181-pBFR1-Bfr1 FL-yeGFP (RJP2203). To generate plasmids carrying multiple mutations (RJP2208; YEplac181-pBFR1-*bfr1mut2A*-yeGFP, RJP2207; YEplac181-pBFR1-*bfr1mut3A*-yeGFP, RJP2206; YEplac181-pBFR1-*bfr1mut4A*-yeGFP, RJP2205; YEplac181-pBFR1-*bfr1mut5A*-yeGFP, RJP2204; YEplac181-pBFR1-*bfr1mut6A*-yeGFP), two-step overlap extension PCR was performed with oligonucleotides RJO6367 and RJO5432 together with mutation specific primers. The corresponding PCR products contained *Kpn*I and *Sac*I restriction sites at their ends. PCR products with lower numbers of mutations were used as templates for generation of additional mutations. The PCR products were then digested with *Kpn*I and *Sac*I and ligated into YEplac181.

Plasmid RJP2209 (carrying a AUG-less *ERG4*) was created as follows. A PCR product of the *ERG4* coding region (lacking the part encoding the first 101 amino acids), together with the 3'UTR, was amplified by PCR with oligonucleotides RJY6527 and RJY6528, introducing *BamH*I and *Hind*III restriction sites. The digested product was ligated into vector pRS316.

**Yeast extracts preparation and subcellular fractionations**

50 $OD_{600}$ units of logarithmically growing yeast cells were harvested and washed once with ice cold water. For subcellular fractionation into membrane and cytoplasmic fractions, we followed a protocol described (Syed et al. 2018). In brief, cells were washed two times with cold potassium phosphate buffer (100 mM potassium phosphate pH 7.5, 1.2 M sucrose) and lysed using glass beads (four pulses of 60 seconds with breaks of 60 seconds on ice) in 500 µl low-salt lysis buffer 1 (20 mM HEPES/KOH pH 7.5, 140 mM potassium acetate, 1 mM magnesium acetate, 1 mM DTT, 250 mM sucrose, 1x EDTA-free Protease Inhibitor Cocktail (Roche)). Cell debris was removed by spinning twice at 800 *x*g for 5 minutes. The resulting whole cell extract (WCE) was diluted to 0.1 µg/µl and 200 µl fractionated at 6,000 *x*g for 5 minutes (M6), 18000 *x*g for 30 minutes (M18) and supernatant (C). Pellets M6 and M18 were resuspended in 200 µl low-salt lysis buffer 1 and 15 µl were used for SDS-PAGE followed by Western blotting. For RNAse A treatment experiments, after removing cell debris and dilution to 0.2 µg/µl, lysates were divided into two equal parts and one part was treated with RNAse A (100 µg/ml) for 20 minutes at room temperature. The other part was considered as a mock treatment.

**Supplementary table ST1.** Yeast strains used in this study.

| Name | Essential genotype | Plasmid(s) | Origin |
|---|---|---|---|
| RJY358 | *MAT**a** ade2-1 trp1-1 can1-100 leu2-3,112 his3-11,15 ura3* | - | |
| RJY925 | *MAT**a**/MAT**alpha**, ade2-1/ade2-1 trp1-1/trp1-1 can1-100/can1-100 leu2-3,112/leu2-3,112 his3-11,15/his3-11,15 ura3* GAL *psi+* | - | |
| RJY4680 | *MAT**a** Bfr1p-yeGFP::HIS3MX6* | - | This study |
| RJY5142 | *MAT**a** Bfr1p(1-397)-yeGFP::KanMX6* | - | This study |
| RJY5204 | *MAT**a** bfr1mut6A-yeGFP::HIS3MX6* | - | This study |
| RJY4626 | *MAT**a** bfr1::HIS3MX6* | - | This study |
| RJY5146 | *MAT**a*** | RJP1870 | This study |
| RJY5148 | *MAT**a** Bfr1p-yeGFP::HIS3MX6* | RJP1870 | This study |
| RJY5417 | *MAT**a** Bfr1p(1-397)-yeGFP::KanMX6* | RJP1870 | This study |
| RJY5419 | *MAT**a** bfr1mut6A-yeGFP::HIS3MX6* | RJP1870 | This study |
| RJY5145 | *MAT**a** bfr1::HIS3MX6* | RJP1870<br>RJP2062 | This study |
| RJY5144 | *MAT**a** bfr1::HIS3MX6* | RJP1870<br>RJP2061 | This study |
| RJY5421 | *MAT**a** bfr1::HIS3MX6* | RJP1870<br>RJP2199 | This study |
| RJY5422 | *MAT**a** bfr1::HIS3MX6* | RJP1870<br>RJP2200 | This study |
| RJY5423 | *MAT**a** bfr1::HIS3MX6* | RJP1870<br>RJP2201 | This study |
| RJY5424 | *MAT**a** bfr1::HIS3MX6* | RJP1870<br>RJP2202 | This study |
| RJY5434 | *MAT**a*** | RJP1809 | This study |
| RJY5435 | *MAT**a** bfr1::HIS3MX6* | RJP1809 | This study |
| RJY5436 | *MAT**a** Bfr1p-yeGFP::HIS3MX6* | RJP1809 | This study |
| RJY5437 | *MAT**a** Bfr1p(1-397)-yeGFP::KanMX6* | RJP1809 | This study |
| RJY5438 | *MAT**a** bfr1mut6A-yeGFP::HIS3MX6* | RJP1809 | This study |
| RJY5425 | *MAT**a** erg6:KanMX6* | - | This study |
| RJY5426 | *MAT**a** erg6::KanMX6 bfr1::HIS3MX6* | RJP2203 | This study |
| RJY5427 | *MAT**a** erg6::KanMX6 bfr1::HIS3MX6* | RJP2204 | This study |
| RJY5428 | *MAT**a** erg6::KanMX6 bfr1::HIS3MX6* | RJP2205 | This study |
| RJY5429 | *MAT**a** erg6::KanMX6 bfr1::HIS3MX6* | RJP2206 | This study |
| RJY5430 | *MAT**a** erg6::KanMX6 bfr1::HIS3MX6* | RJP2207 | This study |
| RJY5431 | *MAT**a** erg6::KanMX6 bfr1::HIS3MX6* | RJP2208 | This study |
| RJY5260 | *MAT**a** ERG4-12xMS2V6::KanMX4* | RJP2121<br>RJP1686 | This study |
| RJY5262 | *MAT**a** bfr1::HIS3MX6 ERG4-12xMS2V6::KanMX4* | RJP2121<br>RJP1686 | This study |
| RJY5432 | *MAT**a** Erg4-yeGFP::klTRP1* | RJP1870 | This study |
| RJY5433 | *MAT**a** Erg4-yeGFP::klTRP bfr1mut6A-yeGFP::HIS3MX6* | RJP1870 | This study |
| RJY5439 | *MAT**a** Erg4-yeGFP::klTRP bfr1mut6A-yeGFP::HIS3MX pep4::KanMX6* | - | This study |
| RJY5440 | *MAT**a** Erg4-yeGFP::klTRP scp160::HIS3MX6* | - | This study |
| RJY4683 | *MAT**a** Bfr1p-6HA::KanMX Scp160-myc9::HIS3MX6* | - | This study |
| RJY3206 | *MAT**a** Khd1-GFP::KanMX4* | - | (Syed et al. 2018) |
| RJY3687 | *MAT**a** RPL16a-TEV-ProtA::HIS3MX6* | - | (Hirschmann et al. 2014) |
| RJY4855 | *MAT**a** RPL16a-TEV-ProtA::HIS3MX bfr1::KanMX4* | - | (Hirschmann et al. 2014) |

| | | | |
|---|---|---|---|
| RJY5441 | *MAT**a** Bfr1p-yeGFP::HIS3MX erg4::KanMX4* | RJP2209 | This study |

**Supplementary table ST2.** Plasmids used in this study.

| Name | Short description | Source |
|---|---|---|
| RJP1870 | pRS415-HDEL-DsRED | (Bevis and Glick 2002) |
| RJP2121 | pET296-YcpLac111-CYC1p-1xMCPNLSSV40-2-xyegfp | (Tutucci et al. 2017) |
| RJP1809 | Edc3-mCh URA3 Cen | (Buchan et al. 2008) |
| RJP1686 | YCp50-SCS2-TMD-2xRFP | (Loewen et al. 2007) |
| RJP2000 | pRS316-pBFR1-Bfr1 FL-yeGFP | This study |
| RJP2199 | pRS316-pBFR1-*bfr1mutR38A*-yeGFP | This study |
| RJP2200 | pRS316-pBFR1-*bfr1mutH79A*-yeGFP | This study |
| RJP2061 | pRS316-pBFR1-*bfr1mutK138A*-yeGFP | This study |
| RJP2201 | pRS316-pBFR1-*bfr1mutF211A*-yeGFP | This study |
| RJP2202 | pRS316-pBFR1-*bfr1mutY225A*-yeGFP | This study |
| RJP2062 | pRS316-pBFR1-*bfr1mutF239A*-yeGFP | This study |
| RJP2209 | pRS316-(-AUG) *ERG4* | This study |
| RJP2203 | YEplac181-pBFR1-Bfr1 FL-yeGFP | This study |
| RJP2204 | YEplac181-pBFR1-*bfr1mut6A*-yeGFP | This study |
| RJP2205 | YEplac181-pBFR1-*bfr1mut5A*-yeGFP | This study |
| RJP2206 | YEplac181-pBFR1-*bfr1mut4A*-yeGFP | This study |
| RJP2207 | YEplac181-pBFR1-*bfr1mut3A*-yeGFP | This study |
| RJP2208 | YEplac181-pBFR1-*bfr1mut2A*-yeGFP | This study |
| RJP148 | pRS316 (URA3, CEN6) | (Janke et al. 2004) |
| RJP143 | YEplac181 (LEU2), 2µm | (Gietz and Schiestl 2007) |

**Supplementary table ST3.** Oligonucleotides used in this study.

| RJO | Name | Sequence (5‘ - 3‘) | Purpose |
|---|---|---|---|
| 5238 | Bfr1_R38A_Fw | GAAATCGGTTTAATTGCCAAGCAAATCGATCAA | Mutation PCR |
| 5239 | Bfr1_R38A_Rw | TTGATCGATTTGCTTGGCAATTAAACCGATTTC | Mutation PCR |
| 5240 | Bfr1_H79A_Fw | CGTAGAAGCAACATTGCCGACTCTATTAAGCAA | Mutation PCR |
| 5241 | Bfr1_H79A_Rw | TTGCTTAATAGAGTCGGCAATGTTGCTTCTACG | Mutation PCR |
| 5242 | Bfr1_K138A_Fw | GAAAAACTACTAGTCGCCGAAATGCAATCTTTG | Mutation PCR |
| 5243 | Bfr1_K138A_Rw | CAAAGATTGCATTTCGGCGACTAGTAGTTTTTC | Mutation PCR |
| 5244 | Bfr1_F211A_Fw | AAAAGACAAACTTTAGCCAACAAACGTGCTGCC | Mutation PCR |
| 5245 | Bfr1_F211A_Rw | GGCAGCACGTTTGTTGGCTAAAGTTTGTCTTTT | Mutation PCR |
| 5246 | Bfr1_Y225A_Fw | AAGCGTGACGAATTAGCCAGTCAAATCAGACAG | Mutation PCR |
| 5247 | Bfr1_Y225A_Rw | CTGTCTGATTTGACTGGCTAATTCGTCACGCTT | Mutation PCR |
| 5248 | Bfr1_F239A_Fw | GACTTTGACAACGAAGCCAAATCATTCAGAGCC | Mutation PCR |
| 5249 | Bfr1_F239A_Rw | GGCTCTGAATGATTTGGCTTCGTTGTCAAAGTC | Mutation PCR |
| 4924 | Bfr1::HA C-terminal S3_Fw | AAAAGATTGAAAGAACAGGAAGAGTCTGAAAAAGATA AAGAAAAATCGTACGCTGCAGGTCGAC | Gene tagging |
| 4925 | Bfr1::HA C-terminal S2_Rw | AATGAAGAAAGATCAGGAGAAAAATTTTTTTCTACTTC AGGTTTAATCGATGAATTCGAGCTCG | Gene tagging |
| 5587 | Eag1 Termi Bfr1_Fw | GGCCGGCCGACCTGAAGTAGAAAAAAATTTTTC | Cloning |
| 5588 | Terminator Bfr1 SacI_Rw | CGAGCTCGGAGGAAAGAATTGGCTGGTAAG | Cloning |
| 6367 | Bfr1 promotor_KpnI_Fw | GGGGTACCGGGCGTAAGAACGAATTTGAAG | Cloning |
| 5432 | yEGFP-Stop-NheI-Rw | GCCGCTAGCTTATTTGTACAATTCATCCATACC | Cloning |
| 5589 | S3_bfr1 1-397AA_Fw | ccaacactaattgctactttggccgaattagacgtaactgtcccaatcCGTA CGCTGCAGGTCGAC | Gene tagging |
| 5590 | S2_bfr11-397AA_Rw | agtaatgaagaaagatcaggagaaaaatttttttctacttcaggtttaATCGA TGAATTCGAGCTCG | Gene tagging |
| 3006 | Bfr1_ko_S1_for | TCAACGTAATAGCATATTTTCTAACAACACAGCCATT GCCCGTACGCTGCAGGTCGAC | Gene deletion |
| 3007 | Bfr1_ko_S2_rev | TGAAGAAAGATCAGGAGAAAAATTTTTTTCTACTTCA GGTATCGATGAATTCGAGCTC | Gene deletion |
| 5828 | ERG4 deltion Fw S1 | CAGATACGGATATTTACGTAGTGTACATAGATTAGCA TCGCTATGCGTACGCTGCAGGTCGAC | Gene deletion |
| 5820 | ERG4 C-term tag Fw S3 | GAGTATTGTAAACATTGCCCTTACGTCTTTATTCCTTA TGTTTTCCGTACGCTGCAGGTCGAC | Gene tagging |
| 5821 | ERG4 C-term tag Rw S2 | ACTGTAAAATAAGTTAATGAAGTGGATAGAAAAAGAA AATAACTA ATCGATGAATTCGAGCTCG | Gene tagging |
| 5826 | ERG6 deltion Fw S1 | AAAAAAACAAGAATAAAATAATAATATAGTAGGCAGC ATAAGATGCGTACGCTGCAGGTCGAC | Gene deletion |
| 5827 | ERG6 deltion Rw S2 | ATATATCGTGCGCTTTATTTGAATCTTATTGATCTAGT GAATTTAATCGATGAATTCGAGCTCG | Gene deletion |
| 6320 | ERG4_NewMS2_F: | ATGAGTATTGTAAACATTGCCCTTACGTCTTTATTCCT TATGTTTTCTAGccgctctagaactagtggat | Gene tagging |
| 6321 | ERG4_NewMS2_R: | ATATACAAACTGTAAAATAAGTTAATGAAGTGGATAG AAAAAGAAAATAAgatatcacctaataacttcgtatag | Gene tagging |
| 6603 | PEP4 -del-S1-Fw | CTAGTATTTAATCCAAATAAAATTCAAACAAAAACCAA AACTAACATGCGTACGCTGCAGGTCGAC | Gene deletion |
| 6604 | PEP4 -del-S2-Rw | TAGATGGCAGAAAAGGATAGGGCGGAGAAGTAAGAA AAGTTTAGCTCAATCGATGAATTCGAGCTCG | Gene deletion |
| 2509 | SCP160_ko_ fw S1 | TAAAATATACTTCCCACACCCCCTCCTTCCATTATAAC TGCACGTACGCTGCAGGTCGAC | Gene deletion |
| 2510 | SCP160_ko_ rev S2 | GCCAAAATCTATATTGAAAAAAATTGGTTTCAAAGAG CTTGTATCGATGAATTCGAGCTC | Gene deletion |
| 6324 | IMH1_qPCR_Fw_Set 2 | ATTGACGACGCAGTGGATAC | qPCR |
| 6325 | IMH1_qPCR_Rw_Set 2 | CTGGAATTTCCGAGTTCTGC | qPCR |

| | | | |
|---|---|---|---|
| 6326 | RUD3_qPCR_Fw_ Set 1 | TTTCGTGTCCATACCCAGAG | qPCR |
| 6327 | RUD3_qPCR_Rw_ Set 1 | CCGGCCTGTTGTTTCTTATC | qPCR |
| 6332 | ERG4_qPCR_Fw_ Set 2 | TGCCATGTGGACCTTGTATG | qRT-PCR |
| 6333 | ERG4_qPCR_Rw_ Set 2 | GTCATGATCCTCCCAAAACG | qRT-PCR |
| 6334 | OSH7_qPCR_Fw_ Set 1 | TGCAGAACAAACGAGTCACC | qPCR |
| 6335 | OSH7_qPCR_Rw_ Set 1 | CCATCATTGCAGCACTTGAG | qPCR |
| 6338 | SGM1_qPCR_Fw_ Set 1 | GGGAAGGGGAATCAATGAAC | qPCR |
| 6339 | SGM1_qPCR_Rw_ Set 1 | GGCCGTTAGAATTCGATAGC | qPCR |
| 2920 | Act1_qPCR_1_Fw | TCAGAGCCCCAGAAGCTTTG | qPCR |
| 2921 | Act1_qPCR_1_Rw | TTGGTCAATACCGGCAGATTC | qPCR |
| 4139 | 18S_qPCR_Fw | TCAACACGGGGAAACTCACC | qRT-PCR |
| 4148 | 18S_qPCR_Rw | CTAAGAACGGCCATGCACCA | qRT-PCR |
| 4488 | MID2_qPCR_Fw | ATGGAGCAAAGCTCCCTTTT | qPCR |
| 4489 | MID2_qPCR_Rw | GCTCTCCACCGCTATTGGTA | qPCR |
| 5568 | Spike-qPCR_Fw | CACACTCATCAGCGACACGA | qPCR |
| 5569 | Spike-qPCR_Rw | CCAAAGCGGTGACTTTCTGC | qPCR |
| 6527 | BamHI_ERG4 _102_Fw | CGCGGATCCATTTGTGCGGAATTTTATCAC | Cloning |
| 6528 | ERG4_3UTR_Hindii i_Rw | CCCAAGCTTGAATTTTCAAAGAATGACTTG | Cloning |
| 3008 | Bfr1_3'UTR_rev | CATGAGGAAAGAATTGGCTGG | Gene tagging |
| 4918 | bfr1_5'UTR_fw | TAACTGATCTCGACGACGTTG | Gene tagging |

**References**


Bevis BJ, Glick BS. 2002. Rapidly maturing variants of the Discosoma red fluorescent protein (DsRed). *Nat Biotechnol* **20**: 83–87.

Buchan JR, Muhlrad D, Parker R. 2008. P bodies promote stress granule assembly in Saccharomyces cerevisiae. *The Journal of Cell Biology* **183**: 441–455.

Gietz RD, Schiestl RH. 2007. High-efficiency yeast transformation using the LiAc/SS carrier DNA/PEG method. *Nat Protoc* **2**: 31–34.

Haim-Vilmovsky L, Gerst JE. 2009. m-TAG: a PCR-based genomic integration method to visualize the localization of specific endogenous mRNAs in vivo in yeast. *Nat Protoc* **4**: 1274–1284.

Hirschmann WD, Westendorf H, Mayer A, Cannarozzi G, Cramer P, Jansen RP. 2014. Scp160p is required for translational efficiency of codon-optimized mRNAs in yeast. *Nucleic Acids Research* **42**: 4043–4055.

Janke C, Magiera MM, Rathfelder N, Taxis C, Reber S, Maekawa H, Moreno-Borchart A, Doenges G, Schwob E, Schiebel E, et al. 2004. A versatile toolbox for PCR-based tagging of yeast genes: new fluorescent proteins, more markers and promoter substitution cassettes. *Yeast* **21**: 947–962.

Loewen CJR, Young BP, Tavassoli S, Levine TP. 2007. Inheritance of cortical ER in yeast is required for normal septin organization. *The Journal of Cell Biology* **179**: 467–483.

Syed MI, Moorthy BT, Jenner A, Fetka I, Jansen R-P. 2018. Signal sequence-independent targeting of MID2 mRNA to the endoplasmic reticulum by the yeast RNA-binding protein Khd1p. *FEBS Letters* **592**: 1870–1881.

Tutucci E, Vera M, Biswas J, Garcia J, Parker R, Singer RH. 2017. An improved MS2 system for accurate reporting of the mRNA life cycle. *Nature Methods* **9**: 777.
